# *In vivo* single-cell ribosome profiling reveals cell-type-specific translational programs during aging

**DOI:** 10.1101/2024.11.02.621639

**Authors:** Clara Duré, Umesh Ghoshdastider, Ramona Weber, Fabiola Valdivia-Francia, Peter F. Renz, Ameya Khandekar, Federica Sella, Katie Hyams, David Taborsky, Merve Yigit, Mark Ormiston, Homare Yamahachi, Mitchell Levesque, Stephanie Ellis, Ataman Sendoel

## Abstract

Somatic stem cells are characterized by their low overall protein synthesis rates, a feature implicated in driving their stemness. However, how aging reshapes the translational landscape of stem cells and how these changes impact their regenerative capacity remains poorly understood. Here, we present an *in vivo* single-cell ribosome profiling strategy to monitor tissue-wide translational landscapes of the young and aged mouse epidermis. By implementing ribosomal elongation-inhibited cell isolation and switching to RNase I ribonuclease for generating ribosomal footprints, we expand the applicability of single-cell ribosome profiling to *in vivo* systems and facilitate the evaluation of triplet periodicity, a hallmark of high-quality ribosome profiling data. Leveraging this strategy and integrating ribosome profiling with single-cell RNA sequencing, we document the *in vivo* translational landscapes of the major epidermal cell types, outline cell-type-specific translational efficiencies and capture heterogeneity in differentiation commitment within stem cell populations. Notably, we identify a pronounced translational reprogramming of AP-1 subunits specifically in aged epidermal stem cells, with functional consequences for keratinocyte behavior. Our study illustrates the power of *in vivo* single-cell ribosome profiling to map cell-type-specific translational programs and offers a scalable strategy for tissue-wide interrogation of translational landscapes at single-cell resolution.

## Introduction

Stem cells are characterized by two features: their ability to self-renew throughout life and to differentiate into other cell types. These functions are closely linked to their precise gene expression regulation. Somatic stem cells exhibit a unique signature marked by high ribosome biogenesis and low protein synthesis rates, a feature that is implicated in driving their stemness independently of cellular proliferation, cell cycle or total mRNA content.^1–5^ The low translation rates of stem cells are largely governed by suppressing mTOR activity, which directly phosphorylates and inactivates the translational repressor eIF4E-binding proteins (4E-BP).^6^ Lower mTOR signaling and protein synthesis rates are associated with enhanced longevity, indicating a protective role of low protein turnover in aging.^7,8^

As stem cells become activated, mTOR signaling acts as an important nexus to coordinate the differentiation program, with subsequent increases in overall protein synthesis rates.^4,9,10^ A tight regulation of the protein synthesis machinery is thereby essential for stem cell function, as demonstrated in hematopoietic stem cells, where both reduced or elevated protein synthesis rates impair engraftment.^2^ However, how exactly the cellular processes during aging shape the translational landscape of stem cells and how these changes impact their regenerative capacity remains poorly understood.

Ribosome profiling enables the monitoring of translational landscapes and has transformed our understanding of post-transcriptional control across diverse cellular contexts.^11–13^ The technique is based on the deep sequencing of ribosome-protected fragments and provides a global snapshot of actively translating ribosomes at single-codon resolution.^12^ Nevertheless, a major drawback of ribosome profiling remained until recently^14,15^ its need for large numbers of cells, limiting its application to study cellular heterogeneity or rare cell types like stem cells in their native environment. The systematic exploration of the translational landscape of stem cells requires an efficient *in vivo* tissue isolation strategy that rapidly blocks ribosomal elongation, coupled with a scalable single-cell ribosome profiling strategy to monitor tissue-wide translation.

Here, we present an *in vivo* single-cell ribosome profiling strategy for the mouse epidermis to comprehensively monitor tissue-wide alterations in the translational landscape during aging. Building on the elegant, recently reported single-cell ribosome profiling protocol^15^, we introduced several critical modifications. Importantly, we implemented a rapid cell isolation process under conditions of ribosomal elongation inhibition and switched to the conventional RNase I-based ribonuclease for generating ribosomal footprints, as commonly utilized in bulk ribosome profiling.^12,16^ Compared to the *Micrococcal Nuclease* (MNase) used in the original protocol, which cleaves following guanine bases^15,17,18^, the standard RNase I lacks any nucleotide bias, resulting in true ribosome-protected fragments of 29-30 nucleotides.^11,12,16^ RNase I-based ribosome profiling therefore facilitates a straightforward evaluation of triplet periodicity, a hallmark feature of high-quality data, as it reflects the ribosome’s movement codon by codon. Consequently, using RNase I eliminates the need for complex machine learning approaches to correct for MNase’s sequence bias, circumventing the risk of overfitting the data. Our study substantially broadens the applicability of single-cell ribosome profiling, offers a scalable strategy for tissue-wide *in vivo* monitoring of translational landscapes at single-cell resolution and uncovers cell-type-specific translational programs in aging.

## Results

### An RNase I-based *in vivo* single-cell ribosome profiling strategy

To develop an *in vivo* single-cell ribosome profiling strategy for the mouse epidermis, we built upon the previously reported single-cell protocol^15^, which leverages a miniature one-pot reaction facilitated by a low-volume liquid handler system. We modified two key aspects. First, to enable efficient *in vivo* translation elongation inhibition, we injected high-dose cycloheximide (CHX) into the mouse tail vein (Figure 1A). The rapid translation elongation inhibition of mouse organs^19^ enables us to prepare epidermal cells by accelerated enzymatic and mechanical isolation and generate single-cell suspensions under continuous submersion in cycloheximide. We then sorted viable epidermal single cells into 384-well plates pre-filled with polysome buffer, where cells are immediately lyzed.

**Figure 1.**
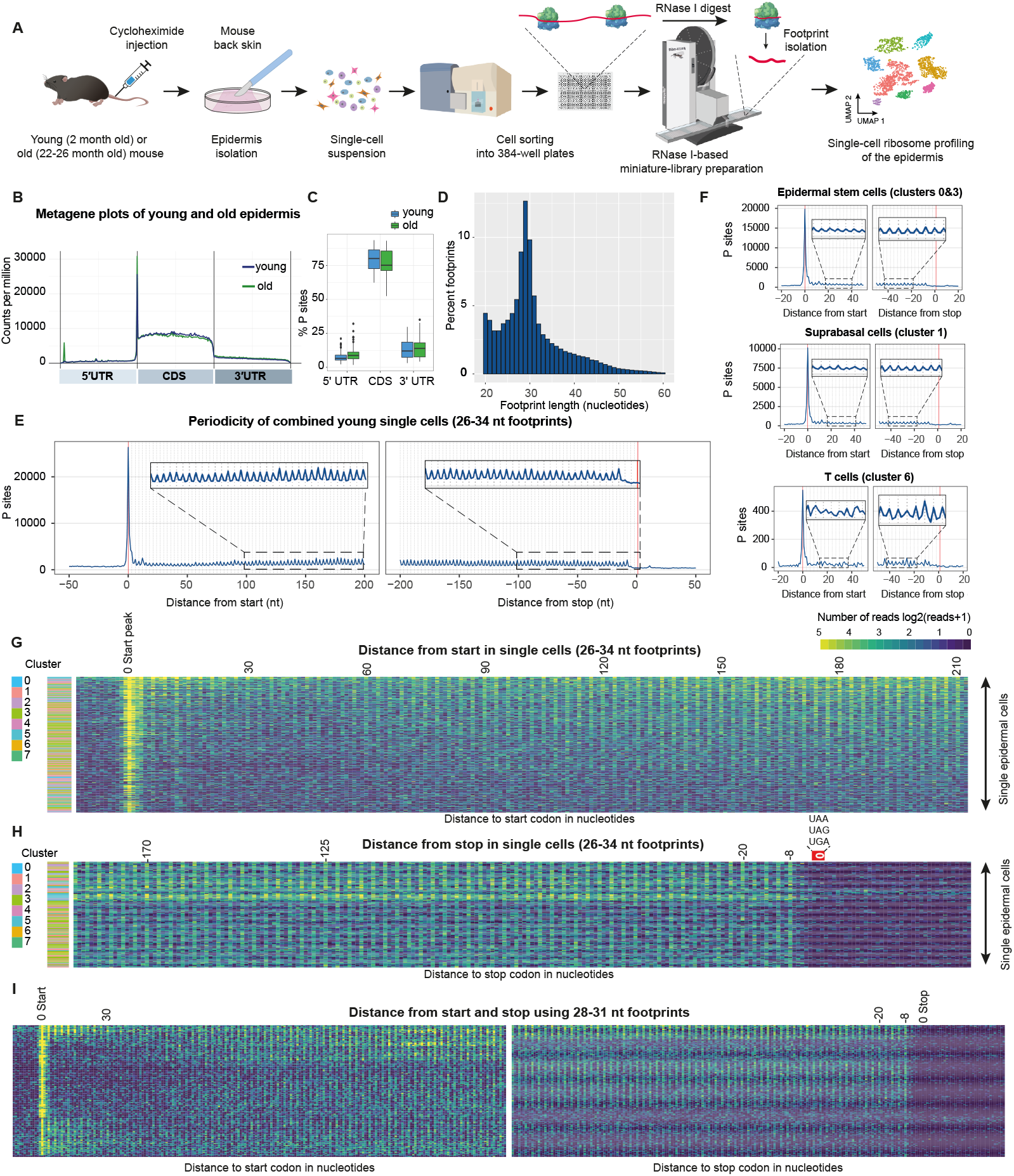
*In vivo* single-cell ribosome profiling of the mouse epidermis. (A) Schematic outline of the RNase I-based *in vivo* single-cell ribosome profiling strategy to monitor the translational landscape of the mouse skin. To efficiently block ribosomal elongation, cycloheximide was injected into the tail vein of female young (2 months) or old (22-26 months) mice before euthanasia. Single-cell suspension was generated via enzymatic and mechanic isolation of the back skin. Single cells were sorted into 384-well plates pre-filled with lysis buffer, followed by an RNase I-based library generation using the SPT Mosquito liquid handler. (B) Metagene analysis of *in vivo* single-cell ribosome profiling of the mouse skin shows footprint enrichment across the coding sequence (CDS). UTR: untranslated region. (C) Distribution of the P site of mapped reads within the 5′UTR, CDS or 3′UTR. (D) Ribosome-protected fragment length distribution shows an enrichment between 28-30 nucleotides, similar to bulk ribosome profiling. The histogram shows the average footprint length distribution of a young epidermis 384-well plate. (E-F) Distinct triplet periodicity of the *in vivo* single-cell ribosome profiling data, the main feature of high-quality ribosome profiling data, is detected when combining all young cells (E) or across the different cell types (F). (G-H) Triplet periodicity can also be detected in single cells *in vivo*. Heatmap showing the coding sequence 200 nucleotides downstream of the start site (G) and the 200 nucleotides upstream of the stop site (H), with position 0 defined as the last nucleotide of the stop codon. The first nucleotide of the P site codon of each footprint is displayed between 26-34 nucleotides (classic range for ribosome profiling gene quantification). Each row represents a single cell. Cluster annotations refer to Figure 2A. (I) Triplet periodicity across the coding sequence is even more distinct when considering only the footprints between 28-31 nucleotides (typical ribosome-protected fragments). Displayed is the P site of each footprint. Each row represents a single cell.

Second, we considered the advantages of conventional RNase I-based ribonuclease treatment for generating ribosomal footprints, as typically utilized in bulk ribosome profiling. To establish an RNase I-based single-cell ribosome profiling protocol, we first adjusted the ribonuclease digestion stop reaction of the original protocol (Methods) and tested the classic RNase I concentrations used to generate bulk ribosome profiling libraries (0.5-2 U/µg RNA). We then generated single-cell ribosome profiling libraries and evaluated the corresponding metagene plots. Surprisingly, neither conventional RNase I nor T1 ribonuclease concentrations, occasionally also used in bulk ribosome profiling^19^, resulted in triplet periodicity or start peak accumulation and instead showed substantial reads in the 3’UTR (Figure S1-AC), a feature of incomplete ribonuclease digestion. We subsequently increased the RNase I and T1 concentrations. Increasing the concentration of ribonucleases up to 10x did not noticeably resolve the reads in the 3’UTR. However, while increasing the RNase I to 100x (50 U/µg RNA) began to decrease the proportion of 3’UTR reads, using 1000x (500 U/µg RNA) of the standard RNase I concentration generated a start codon peak and remarkably clean 3’UTRs (Figure S1A). Surprised by the necessity for such high RNase I concentrations, we evaluated the 1000x concentration by comparing it to the standard units per RNA input (U/µg RNA) and the unit concentration in the lysate (U/µl). We found that the ribonuclease concentration used in bulk ribosome profiling closely aligns with the 1000x conditions (0.2 U/µl vs. 0.05-0.1 U/µl used in bulk) applied in our 50 nl one-pot reaction. These results indicate that low-input ribosome profiling samples should prioritize overall enzyme concentration as opposed to relying on enzyme units per RNA input.

Using our optimized RNase I-based *in vivo* ribosome profiling strategy, we then set out to systematically monitor the translational landscape of the mouse epidermis. Several lines of evidence indicate that our single-cell ribosome profiling data represent *bona fide* translation. First, similar to bulk ribosome profiling, we observed clearly reduced fractions of 5′UTR and 3’UTR reads compared to total RNA sequencing, reflecting translation across the coding sequence, with varying levels of upstream and downstream open reading frames^13^ (Figure 1B-C). Second, employing RNase I, we found a fragment length distribution comparable to bulk ribosome profiling, with a distinct peak at 29 nucleotides (Figure 1D). The main ribosome-protected fragment lengths of 28-30 nucleotides showed the classic P site offset (distance from 5′ end to P site) of 12 nucleotides (Figures S1D). Third, our data showed a strong start peak, reflecting the efficient inhibition of translation elongation by cycloheximide but not 40S scanning and translation initiation during the single-cell suspension preparation (Figure 1B-F, S2D). Fourth, by examining the hallmark feature indicative of true translation, we observed an excellent triplet periodicity throughout the entire coding sequence (Figure 1E-F). This was also evident when evaluating diverse cell types, including stem cell populations or skin-resident immune cells such as T cells (Figure 1F, S2C, S3C). Importantly, when zooming into single cells, detected strong triplet periodicity within single epidermal cells, considering the footprint lengths ranging from 26-34 nucleotides (Figure 1G-H).

As expected, we observed the most defined triplet periodicity when considering only the classic footprint lengths of 28-31 nucleotides, which yielded more sparse but even cleaner periodicity patterns in the heatmaps (Figure 1I, S2A). Furthermore, we also found excellent periodicity in single cells upstream of the stop codon, underscoring *bona fide* translation across the complete coding sequence (Figure 1H-I, S2A, S2D). Collectively, the assessment of the ribosome profiling quality suggests that our RNase I-based *in vivo* approach is ready to investigate tissue-wide *in vivo* translational landscapes (Methods).

### Cell-type-specific translational landscapes of the young epidermis

To determine translational changes in aging, we next sought to monitor the translational landscapes of young (2 months) and old (22-26 months) mouse epidermis, following the distinct morphological changes that occur in the skin during aging (Figure 2B-E). After rRNA removal and filtering out cells with low unique molecular identifier (UMI, cutoff of 100) counts, we retained 1068 young and 939 old epidermal cells (Figure 2A). Using Uniform Manifold Approximation and Projection (UMAP) and cell-type specific marker genes, we could identify the main cell types present in the epidermis, including epidermal stem cells, suprabasal cells, bulge stem cells, upper hair follicle cells, T cells, melanocytes as well as macrophages and dendritic cells (Figure 2A, F-G, S3A-B). We first focused on the young epidermis. To showcase the coverage and cells uniquely translating marker genes, we selected 9 single cells, which exhibited excellent coverage across the whole coding sequence of cluster-defining marker genes. Cells 1 and 2 predominantly translated *Krt14*, an EpSC marker, clearly distinct from the differentiated suprabasal cells 5 and 6, translating *Krt10*. Cells 3 and 4 expressed *Krt14* and G2/M markers such as *Top2a*. Furthermore, cells 7 and 8 translated the upper hair follicle marker *Krt79*, one residing in the basal layer (*Krt14*-positive) and the other in the suprabasal layer (*Krt10*-positive). The macrophage in track 9 specifically exhibited footprints in *Cd74*. The housekeeping gene *Actb* was ubiquitously translated across these 9 cells (Figure 2I). Collectively, these results highlight the sensitivity and specificity of our established *in vivo* single-cell ribosome profiling approach.

**Figure 2.**
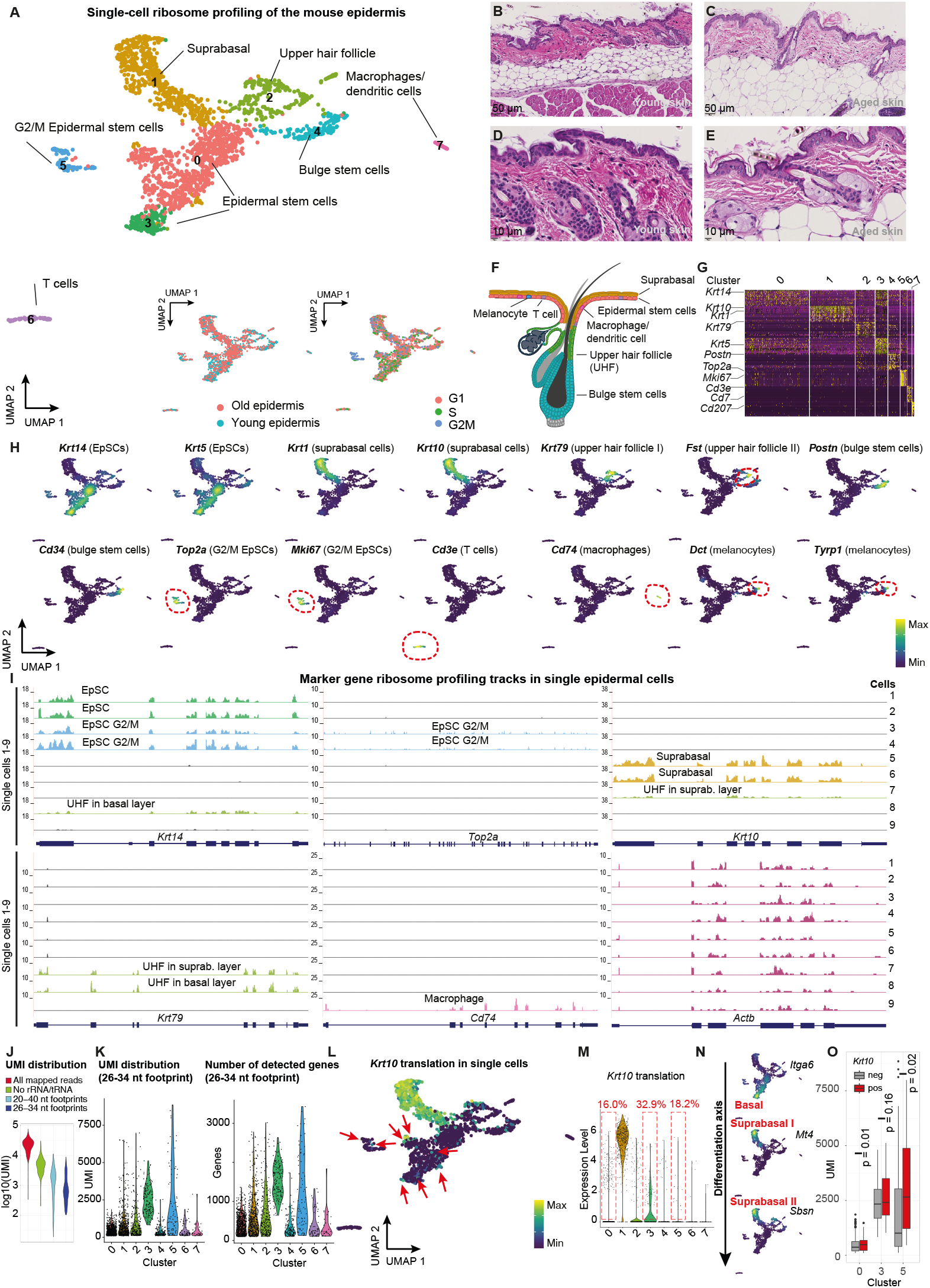
Translational landscape of the young and old epidermis. (A) Uniform manifold approximation and projection (UMAP) of 2,007 epidermal cells identifies 8 cell types based on their translation programs. The UMAP incorporates data from 1068 epidermal cells from three young animals (2 months) and 939 cells from three old animals (22-26 months). Insets show the contribution of young and old cells (left panel) and the cell cycle stage (right panel). (B-E) Hematoxylin and eosin (H&E) staining of young (B, D, 2 months) and aged (C, E, 25 months) mouse skin. Scale bars, 50 µm (B, C) and 10 µm (D, E). (F) Schematic representation of the anatomical localization of different cell populations in the adult mouse epidermis. (G) Heatmap displaying the single-cell ribosome profiling expression profiles of the top marker genes across the young and old epidermis. (H) Nebulosa density plots visualize the expression of cell-type-specific marker genes across the UMAP. (I) Marker gene ribosome profiling tracks in single epidermal cells show translation of *Krt14, Top2a, Krt10, Krt79, Cd74* and *Actb*. (J) Unique molecular identifier (UMI)-corrected unique reads distribution for all aligned reads, following rRNA and tRNA removal, when considering the footprints between 20-40 nucleotides and the final filtered cells with a UMI cutoff of 100. Displayed is the example of a 384-well plate with cells from a young mouse. (K) Unique read count distribution and number of detected genes when considering the stringent cutoff of 26-34 nucleotides for footprints. Clusters refer to Figure 2A. (L) Single-cell expression profiles (as opposed to Nebulosa density plots in Figure 2H) show *Krt10* translation in single differentiation-committed epidermal stem cells. (M) *Krt10* expression in the single-cell ribosome profiling data of the mouse skin reveals 32.9% *Krt10-*positive epidermal stem cells in cluster 3. Clusters refer to Figure 2A. (N) The general differentiation axis is highlighted by the translation of the marker genes *Itga6* (epidermal stem cells), *Mt4* (early suprabasal) and *Sbsn* (late suprabasal). (O) Unique read count distribution of *Krt10*-positive and -negative clusters show significantly higher read counts in clusters 0 and 5. P values indicate a Wilcoxon test.

Overall, after genome alignment, we observed a duplicate-corrected UMI distribution with a median of 22,252 unique reads per cell (Figure 2J). However, rRNA and tRNA filtering, which constitute 75-80% of total reads, reduced the median UMI to 4,653 (Figure 2J). Due to the one-pot reaction used in the single-cell ribosome profiling protocol, which omits the footprint excision step, the footprint length distribution was broader than in bulk ribosome profiling (Figure 1D). Thus, as expected, applying the stringent footprint cutoff of 26-34 nucleotides to ensure accurate translation quantification, the median decreased further to 684 unique reads in young cells (Figure 2J). Generally, we observed cell-type-specific differences in the distribution of unique reads (Figure 2K). The stem cell population, such as epidermal stem cells (cluster 0, median of 500 unique reads) and bulge stem cells (cluster 4, median of 349 unique reads), exhibited low read counts compared to for example differentiated suprabasal cells (cluster 1, median of 810 unique reads). These differences were also evident when assessing the cell-type-specific differences in the triplet periodicity heatmaps, with a sparse signal in single epidermal stem cells of cluster 0, despite the clear triplet periodicity when aggregating all epidermal stem cells (Figures S2C, 1F). Interestingly, a subpopulation of epidermal stem cells (cluster 3, median unique reads of 2583) and epidermal stem cells in the G2/M cell cycle phases (cluster 5, median unique reads of 1834) exhibited high read counts. This difference was not due to higher sequencing saturation in clusters 3 and 5 (Figure S3D). The relative contribution of cluster 3 within epidermal stem cells was diminished in aged skin (0.31 in young vs. 0.067 in old).

We reasoned that, beyond potential technical factors, unique read counts on the same plate capture the differences in translation rates across different cell types. Somatic stem cells, such as epidermal stem cells or bulge stem cells, are known to exhibit low overall protein synthesis rates.^1,3,20^ Our read count observations thus suggested that the stem cell populations in the epidermis exhibit lower translation rates than differentiated cells (Figure 2K). Using O-propargyl-puromycin (OPP) incorporation and immunofluorescence of the mouse skin, we confirmed that suprabasal cells overall exhibited 1.75 times higher protein synthesis rates compared to epidermal stem cells (matching the median UMI difference of 810 vs. 500 in suprabasal vs. epidermal stem cells), in line with low unique read counts representing low translation rates of epidermal stem cells (Figure S4A).

The epidermal stem cell compartment is known to encompass a heterogeneous subpopulation, with *Krt10* as one of the earliest markers for differentiation-committed stem cells, preceding delamination and downregulation of epidermal stemness genes.^21^ Delaminating cells therefore transition through a *Krt10*-positive state, retaining a limited capacity for cell division.^21^ We therefore examined the potential stem cell commitment within the epidermal stem cell population in more detail. In line with previous reports^21^, we found that 32.9% of cluster 3 and 16.0% of cluster 0 epidermal stem cells were *Krt10*-positive, indicating that they not only store *Krt10* mRNA but actively synthesize KRT10 protein in epidermal stem cells (Figure 2L-M). A small fraction of *Krt10*-positive epidermal stem cells remained actively proliferating, with 18.2% of the G2/M epidermal stem cell cluster translating *Krt10* (Figure 2M). Collectively, these results indicate that a subset of epidermal stem cells is committed to differentiation, with active KRT10 translation, while retaining a capacity for cell division.

Previous studies in the hair follicle have shown that increasing translation rates correlated with stem cell commitment and differentiation rather than proliferation.^1^ We then asked whether *Krt10*-positive differentiation-committed epidermal stem cells exhibited altered unique read count distribution compared to *Krt10*-negative cells. While *Krt10*-positive cells in epidermal stem cell cluster 3 did not exhibit a significant difference in read counts, *Krt10*-positive cells in clusters 0 and 5 (G2M) showed significantly increased unique read counts as well as a higher number of detected genes (Figure 2O, S4F). We observed no concomitant increase in *Krt5* or *Krt14* marker gene reads in *Krt10*-positive cells, indicating that *Krt10* detection is not due to a general increase in read counts (Figure S4G). These observations suggest that stem cell commitment at least partly correlates with increased unique read counts, in line with a boost in translation rates during differentiation commitment. Collectively, our findings reveal heterogeneity in the epidermal stem cell population, with subpopulations exhibiting differences in both differentiation commitment and translation rates.

### *In vivo* translational efficiencies during differentiation

Next, we examined cell-type-specific translational efficiencies (TE, defined by ribosome profiling reads/ RNA sequencing reads). Similar to the single-cell ribosome profiling experiments, we collected the epidermis of young (2 months) and aged mice (22-26 months) and generated integrated parallel single-cell RNA sequencing libraries. After stringent filtering, we identified a total of 15,873 cells representing the major cell types of the epidermis (Figure 3A-C, S4B-D). We then explored the translational efficiencies of each cell type by determining the reads in the coding sequence in single-cell ribosome profiling divided by the transcript reads in single-cell RNA sequencing (Methods).

**Figure 3:**
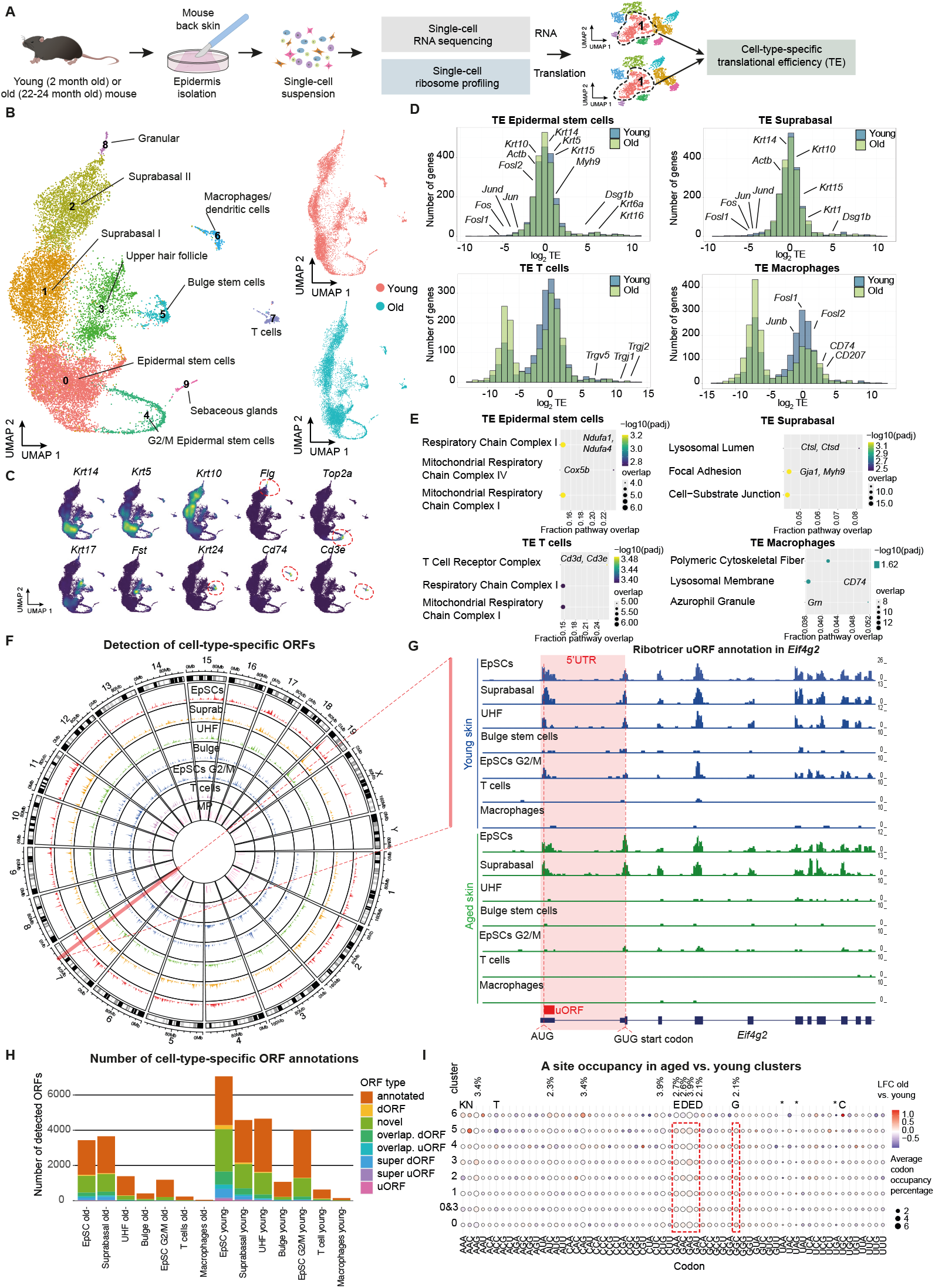
Cell-type-specific translation efficiencies and ORF detection. (A) Schematic representation of the parallel single-cell RNA sequencing and ribosome profiling strategy to monitor cell-type-specific translational efficiencies in the mouse epidermis. (B) Single-cell RNA sequencing and corresponding UMAP of young (2 months) and old (24 months) mouse skin (n = 1 animal per condition). Insets show the contribution of young and old cells. Young = 7426 cells, old = 8447 cells. (C) Nebulosa density plots visualize cell-type-specific marker gene expression. (D) Histogram showing cell-type-specific translational efficiencies (TE) in young epidermal stem cells, suprabasal cells, T cells, and macrophages. TE is calculated as the ratio of CPM from ribosome profiling to CPM from RNA sequencing for each cell type using a pseudobulk approach. A total of 1958 genes with an average TPM/CPM greater than 100 in both RNA and ribosome profiling were included across all cell types. Highlighted genes refer to TE in the young epidermis. (E) GO terms (GO cellular component 2023) enriched in the top 250 genes with the highest translational efficiency (TE) across each of the four cell types in the young epidermis. Representative genes from each category are highlighted. (F) Circos plot illustrating cell-type-specific open reading frames (ORFs) across mouse chromosomes, as identified through Ribotricer and single-cell ribosome profiling data. Only female mice were included in this study. (G) Ribosome profiling tracks of the different epidermal cell types in young and old skin show a strong AUG start peak for the uORF in *Eif4g2*. ORFs were annotated using Ribotricer. (H) Number of cell-type-specific ORF annotations predicted by Ribotricer using the single-cell ribosome profiling data of the mouse epidermis. dORF, downstream ORF. Overlap., overlapping. Super refers to upstream or downstream ORFs that do not overlap with the CDS of the same gene. (I) A site codon occupancy in aged vs. young epidermal cell clusters. Clusters refer to Figure 2A. Reported mouse codon usages are shown for some codons above the clusters. Dot size refers to the average codon occupancy of young and old epidermal clusters.

Zooming into the top TE genes in the young epidermis, we observed that the mitochondrial chain complex I and IV genes were enriched among the most efficiently translated genes in epidermal stem cells (Figure 3D-E). This was in contrast to suprabasal cells, which exhibited high TE of focal adhesion and lysosomal lumen genes. Furthermore, T cells showed high TE in T cell receptor genes and macrophages in granule and lysosomal membrane genes such as *Cd74* (Figure 3D-E). These variations in TE may suggest a link between the translation efficiency of gene cohorts and their relative importance in supporting cell type functions. For example, intestinal stem cells are known to display high mitochondrial activity, which is critical for their function.^22^

On the other hand, lysosomal lumen genes such as cathepsins in suprabasal cells regulate the crosslinking of the cornified envelope proteins involucrin and loricrin during epidermal differentiation^23^, which may be, together with focal adhesions, particularly important for differentiating cells. Of note, the AP-1 subunits, including *Jun, Junb, Jund, Fos, Fosb, Fosl2 or Atf3*, were among the lowest TE genes in epidermal stem cells and suprabasal cells, indicating translational repression of these genes in the interfollicular epidermis (Figure 3D, Table 1).

The classic cell-type-defining marker genes such as *Krt14* and *Krt5* for epidermal stem cells or *Krt1* and *Krt10* for suprabasal cells were translated with overall average TE (Figure 3D). Nevertheless, we observed notable TE switches in the marker genes during differentiation. *Krt10* TE switched from inefficient translation in epidermal stem cells (TE log_2_: -0.61) to more efficient translation in suprabasal cells (TE log_2_: 0.52), as did *Krt1* (from TE log_2_ of 0.31 to 1.86).

The opposite was true for *Krt14*, switching from more efficient translation (TE log_2_: 0.75) in epidermal stem cells to lower translational efficiency in suprabasal (TE log_2_: -0.20), suggesting additional posttranscriptional regulation of these marker genes beyond their transcriptional control.

Expanding beyond cell-type-specific translational efficiencies, we also explored the utility of *in vivo* single-cell ribosome profiling for cell-type-specific annotation of open reading frames (ORFs). A substantial fraction of the cellular proteome, often referred to as the dark proteome, is translated from noncanonical ORFs, many of which encode small ORFs.^24,25^ To systematically explore cell-type-specific ORFs in the skin, we ran Ribotricer, a robust method for detecting actively translating ORFs by assessing the triplet periodicity^26^, to annotate different categories of ORFs (Figure 3H). The identified number of ORFs largely reflected the number of cells and read counts in the different cell types, suggesting that the approach is scalable to increase ORF detection in specific cell types (Figure 3F-H, Table 2). The annotated ORFs were found across the mouse genome, except the Y chromosome, as only female mice were included in the study (Figure 3F). For example, Ribotricer predicted an upstream open reading frame in *Eif4g2*, which was clearly visible also in the footprint tracks, with a pronounced ribosome profiling peak at the predicted 5′UTR uORF start codon (Figure 3G).

Our approach also allowed us to compute codon occupancy at the E, P and A sites of the ribosome (Figures 3I, S4H-I, Methods). Potential differences in A site occupancy in aging can reflect tRNA availability, as tRNA scarcity can lead to empty A sites and higher signals. For example, we observed that two aspartate (D) and two glutamate (E) codons exhibited consistently higher occupancy in the aged epidermis, indicating a potential depletion of these aminoacylated tRNAs in aging (Figure 3I). Glutamate is a major source of the tricarboxylic acid cycle (TCA) cycle intermediate α-ketoglutarate, which serves as a substrate for dioxygenases required for hypoxia signaling and epigenetic networks. Additionally, changes in the extracellular matrix have been shown to trigger metabolic crosstalk between aspartate and glutamate^27,28^, highlighting potential metabolic shifts in the aged epidermis.

### The translational landscape of aging

To investigate the changes in the translational landscape of the aged epidermis, we next integrated the parallel single-cell ribosome profiling and RNA sequencing experiments conducted in young and aged mice (Figure 4B). By contrasting single-cell ribosome profiling from single-cell RNA-sequencing, we reasoned that this strategy would enable us to compute tissue-wide *in vivo* translational efficiencies (TE) changes in aging for each epidermal cell type. We took three different approaches. First, we employed glmGamPoi, an adaptation of the robust DESeq2 pipeline to fit the Gamma-Poisson distribution to single-cell data.^29^ Second, we compared this approach to the standard Seurat pipeline for differential gene expression calculation in single cells.^30^ Third, to independently validate cell-type-specific *in vivo* translational efficiencies, we also directly computed TE from aggregated pseudobulk data for each cell typeindividually, with the advantage of directly calculating p values for TE changes.

**Figure 4:**
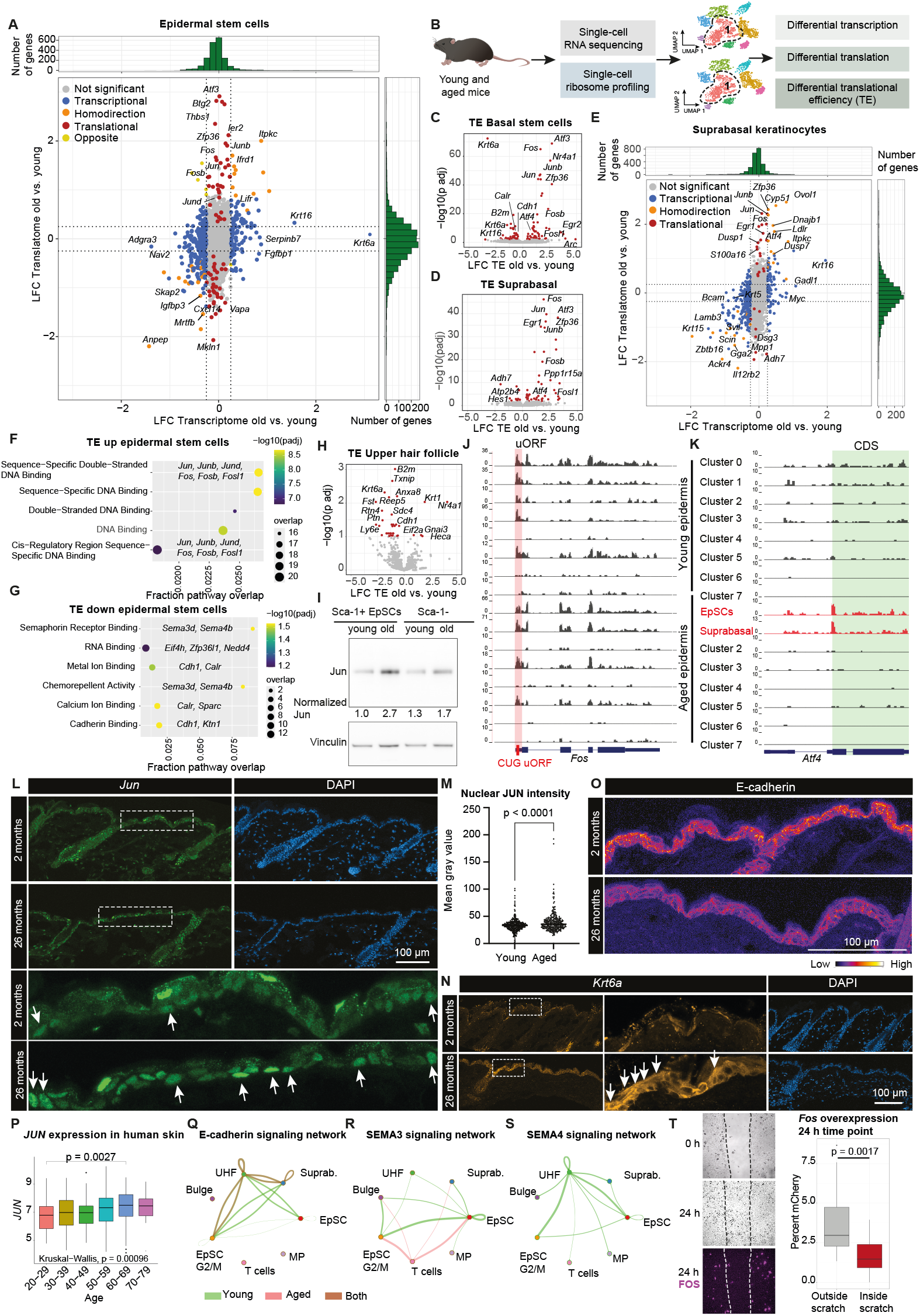
Cell-type-specific alterations in translation efficiencies of the aged epidermis. (A) Scatterplot of transcriptional and translational changes in old versus young epidermal stem cells. Single-cell RNA sequencing and single-cell ribosome profiling data were integrated for the same cluster using glmGamPoi. LFC, log_2_ fold change. Transcriptional changes are in blue (single-cell RNA-seq p adjusted < 0.05 and I LFC I > 0.25), homodirectional changes in orange (single-cell RNA-seq and single-cell ribosome profiling p adjusted < 0.05 and I LFC I > 0.25) and translational changes are in red (single-cell ribosome profiling p adjusted < 0.05 and I LFC I > 0.25). Histograms show the distribution of the changes. (B) Schematic representation of the strategy to combine the *in vivo* single-cell RNA sequencing with the *in vivo* single-cell ribosome profiling to analyze differential transcription, differential translation or translational efficiencies of the different cell types of the aging epidermis. (C-D) Volcano plot of translational efficiency (TE) changes in aged epidermal stem cells or suprabasal cells. Cell-type-specific translational efficiencies were directly computed by a pseudobulk approach using DESeq2 for each cell type. (E) Scatterplot of transcriptional and translational changes in old versus young suprabasal cells. (F-G) Pathways enriched in significantly TE upregulated or TE downregulated genes in old epidermal stem cells. Pathways, BioPlanet 2019. (H) Volcano plot of translational efficiency (TE) changes in aged upper hair follicle cells. Cell-type-specific translational efficiencies were directly computed by a pseudobulk approach using DESeq2. (I) Western blot analysis of cell lysates from young (2 months) and old (26 months) animals. Sca-1+ and Sca-1-epidermal stem cells from both young and old animals were analyzed for JUN expression. Cells were sorted via fluorescence-activated cell sorting (FACS) using the Ly-6A/E (Sca-1) monoclonal antibody and VINCULIN was used as the loading control. Quantifications represent the normalized average of two biological replicates (2 young and 2 old animals), analyzed using ImageJ. (J) Ribosome profiling tracks of *Fos* reveal a uORF at the CUG start codon, translated in several clusters, including epidermal stem cells and suprabasal cells. (K) Ribosome profiling tracks of *Atf4* reveal cell-type-specific start peak induction, specifically in aged epidermal stem cells and suprabasal cells. (L) Immunofluorescence of young (2 months) and old (26 months) back skin shows induction of nuclear JUN expression in aged epidermal stem cells. Lower panels provide a magnified view of the boxed region, arrows indicate nuclear JUN. Scale bars, 100 µm. (M) Quantification of nuclear JUN intensities of young and old epidermal stem cells. 353 young and 282 old cells were quantified. p value indicates an unpaired two-tailed t test. (N) KRT6A is induced in the aged interfollicular epidermis. Magnified views of the boxed KRT6A region are displayed in the middle panels. Arrows indicate KRT6A induction in suprabasal basal. (O) E-cadherin signal is reduced in aged epidermal stem cells, as shown by heatmaps of E-cadherin in young and aged epidermis. (P) *JUN* expression correlates with age in the human skin (Genotype-Tissue Expression (GTEx) portal data). p value indicates a Wilcoxon test. Additionally, a Kruskal-Wallis test was performed. N = 40, 43, 96, 193, 174, 10 (from young to old). (Q-S) Ligand-receptor analysis of young and aged epidermis employing CellChat reveals tissue-wide alterations of E-cadherin and semaphorin signaling. The arrows indicate directional signaling between different cell types and the thickness of the arrow indicates the signaling strength. No semaphorin 4 signaling was detected in the aged epidermis. EpSC, epidermal stem cells. UHF, upper hair follicle. MP, macrophages. (T) *Fos* overexpression impairs keratinocyte migration into the *in vitro* wound. *Fos::P2A::mCherry-*overexpressing and wild-type keratinocytes were seeded into a culture insert and migration into the cell-free gap was tracked over 24 hours with hourly imaging. N = 16 quantified tiles (24-hour time point), from 6 independent experiments. P value indicates a t test.

By coupling the single-cell RNA sequencing and ribosome profiling data, we first focused on the largest cluster and contrasted genome-wide transcriptional and translational changes of aged epidermal stem cells. Some genes, such as *Krt6a* and *Krt16*, were predominantly altered at the transcriptional level, without any concomitant change at the translational level (Figure 4A). *Krt6* and *Krt16* are known to be jointly regulated during epidermal barrier stress, together with a network of genes involved in skin barrier protection and innate immune responses.^31^ This transcription induction does, however, not result in significantly higher translation of *Krt6* and *Krt16* in epidermal stem cells. Other genes, such as *Itpkc, Anpep* or *Ifrd1* exhibited homodirectional changes, with simultaneous alteration at the level of transcription and translation. Finally, we defined a class of genes altered only at the translational level, without concomitant changes in transcription. Among these translationally regulated genes, we observed a noticeable upregulation of AP-1 subunits, including *Jun, Junb, Jund, Fos, Fosb* and *Atf3* (Figure 4A, Table 3). These subunits showed an increase in translation in the absence of any transcriptional induction in aged epidermal stem cells. We obtained similar overall distributions using the Seurat pipeline (Figure S4E).

The induction in translational efficiency of AP-1 subunits was confined to the interfollicular epidermis as none of the other cell types showed alterations of AP-1 subunits (Table 3). Using an independent and robust aggregated pseudobulk approach for each cell type, we also directly computed TE changes with the corresponding p values employing DESeq2^32^. Direct TE calculations confirmed the significant TE changes of AP-1 subunits specifically in epidermal stem cells and suprabasal cells (Figure 4C-D, Table 4). The strong enrichment of AP-1 subunits was also reflected in the enriched gene ontology terms of the TE-high genes, which were dominated by the sequence-specific DNA binding of AP-1 subunits (Figure 4F-G). Of note, AP-1 subunits were among the most translationally repressed genes in young epidermal stem cells (Figure 2D), suggesting a partial derepression in aging. In contrast, upper hair follicle cells, for example, exhibited no AP-1 TE changes and instead revealed a different cohort of genes with alterations in TE (Figure 4H, S4J). These observations argue against the possibility of artifacts arising from AP-1 induction through a stress response during single-cell suspension preparations^33,34^, which are predominantly a transcriptional induction and should also uniformly affect the other cell types, a pattern not evident in our data.

AP-1 changes in translational efficiency during aging were largely maintained during differentiation as the AP-1 subunits were also translationally increased in suprabasal cells (Figure 4D-E). However, while *Krt6a* and *Krt16* exhibited transcriptional induction but strongly reduced TE in epidermal stem cells (Figure 4C), TE of *Krt6a* and *Krt16* was no longer downregulated in suprabasal cells (Figure 4D). These findings suggest that - while *Krt6* and *Krt16* mRNAs are transcriptionally induced in aged epidermal stem cells - this mRNA induction is mainly translated in differentiated cells as opposed to epidermal stem cells, highlighting the power of examining cell-type-specific gene expression cascades along a differentiation axis *in vivo*.

Interestingly, the 5′UTR of the AP-1 subunit *Fos* revealed an obvious uORF, with a pronounced ribosome profiling peak at the CUG start codon (Figure 4J). Similarly, *Jun* 5′UTR is nearly one kilobase in length and regulated by its uORFs^35,36^, which are also highly translated in epidermal stem cells (Figure S4K). Posttranscriptional control has also been reported for other AP-1 subunits.^36^ Thus, these observations suggest a general translational reprogramming of AP-1 subunits and raise the possibility of involving uORF-dependent mechanisms in the control of AP-1 translation. The arguably best-studied uORF-regulated gene, the mammalian Activating Transcription Factor 4 (*Atf4*), is controlled by two uORFs, which repress the translation of *Atf4* in homeostasis.^37,38^ Upon stress, the inhibitory effect of the overlapping uORF is bypassed, resulting in translational ATF4 induction.^39^ Focusing on *Atf4* as a readout for uORF-regulated translational control, we found that it is translationally upregulated in aged epidermal stem cells and suprabasal cells but not other cell types (Figure 4C-E). The induction of the *Atf4* main ORF translation was also clearly detectable in the cell-type-specific ribosome profiling tracks, with a pronounced start site peak in the interfollicular aged clusters (Figure 4K), indicating uORF-controlled translational induction. In summary, these findings suggest that alterations in the translational machinery in aging drive cell-type-specific translational reprogramming, which in part involve uORF-mediated translational control.

To validate the increased translation of AP-1 in aging, we used two orthogonal approaches. We first isolated SCA-1-positive (enriched for epidermal stem cells) and SCA-1-negative epidermal cells by fluorescence-activated cell sorting (FACS) and quantified JUN protein levels by western blot. We found that SCA-1-positive aged epidermal stem cells exhibited a marked upregulation of JUN protein levels (Figure 4I). Second, we evaluated JUN expression in aged skin by immunofluorescence (Figure 4L). While the young skin predominantly showed JUN expression in different hair follicle populations, we observed a pronounced nuclear JUN induction in aged epidermal stem cells, evident also by the higher intensity of nuclear JUN in aged epidermal stem cells (Figure 4M). Furthermore, using the ChIP Enrichment Analysis (ChEA) database, we identified a strong enrichment of *Jun* and *Jund*-regulated genes among the transcripts upregulated in aged epidermal stem cells (Figure S4M), in line with higher AP-1 levels. Moreover, higher *JUN* expression in the human skin also correlated with age (Figure 4P). Consistent with our findings of altered *Krt6a* translation in the aged suprabasal layer, we also observed an induction of KRT6A in immunofluorescence sections of aged epithelia, with a pronounced signal in suprabasal layers (Figure 4N). Collectively, these data validate the single-cell ribosome profiling findings and indicate nuclear AP-1 subunit induction specifically in the aged interfollicular epidermis.

We also observed TE reductions in aged epidermal stem cells of mRNAs that play a critical role in epidermal homeostasis, such as E-cadherin (*Cdh1*) expressed in adherens junctions (Table 4). We validated these findings by immunofluorescence sections and observed a less distinct E-cadherin signal in aged epidermal stem cells (Figure 4O). Furthermore, semaphorin signaling in epidermal stem cells similarly showed lower TE (Figure 4G). Semaphorins are known to have immunomodulatory effects on T lymphocytes, dendritic cells and macrophages and decreased semaphorin 3 levels have been observed in autoimmune diseases such as psoriasis.^40,41^ Probing ligand-receptor interactions by leveraging our single-cell ribosome profiling data and CellChat^42^, we observed that the E-cadherin and semaphorin signaling alterations have tissue-wide consequences for the cell-cell signaling network, including re-routing of cadherin signaling away from aged epidermal stem cells and increased semaphorin 3 signaling from aged epidermal stem cells to T cells (Figures 4Q-S, S4L). Moreover, we also observed a TE reduction of calreticulin (*Calr*) and *B2m* (Figure 4C-D, G), which have been recently shown to serve as “eat-me” and “don’t-eat me” signals on hematopoietic stem cells. These signals play a critical role in shaping hematopoietic clonality^43,44^, raising the possibility of also influencing clonal expansions in aged epithelia.^45–47^ Together, these findings suggest that the specific TE alterations in aged epidermal stem cells could directly impact a wide range of epidermal functions and may underlie the declined skin function observed in aged epidermis.

Finally, we examined whether translational reprogramming of AP-1 could alter keratinocyte function. A hallmark of the aged epidermis and stem cell decline is the reduced capacity for wound healing.^48^ To assess the possibility that translational reprogramming of AP-1 may alter keratinocyte function in aging, we overexpressed the AP-1 subunit *Fos* in keratinocytes and performed epithelial scratch assays as a measure for their wound healing capacity. We found that AP-1 overexpression impaired keratinocyte migration into the *in vitro* wound (Figure 4T), suggesting that translational reprogramming of AP-1 in aging may alter keratinocyte function.

## Discussion

Translation is the most energy-demanding process in a cell. Thus, its precise regulation is critical to conserve cellular energy and ensure that only necessary transcripts are translated to support cellular functions.^4,49^ Here, we develop an *in vivo* single-cell ribosome profiling strategy for the mouse epidermis to monitor tissue-wide translational landscapes. Importantly, we couple *in vivo* ribosomal elongation blocking, efficient single-cell preparation and a critical switch to employing an RNase I-based ribonuclease foot-printing method. Our strategy enables the direct monitoring of triplet periodicity, without the need for a machine learning strategy to correct for the sequence bias of the MNase. We show that our approach results in excellent triplet periodicity across diverse cell types, ranging from epidermal stem cells to T cells, a hallmark feature of high-quality ribosome profiling data. It is worth mentioning that our sequencing saturation is currently only at 15-25% (Figure S1F), which could be substantially increased depending on biological questions, the necessary number of quantified genes and budget considerations. Additionally, implementing rRNA removal into the single-cell ribosome profiling workflow could, in the future, strongly enhance the yield of unique reads per cell (Figure 2J). Therefore, the described *in vivo* approach is scalable and readily adaptable to other tissues and organisms, and, as such, provides a general scheme for quantifying mRNA translation and translational efficiencies at single-cell resolution across tissues.

We leverage our outlined strategy to document the translational landscape of the young and aged epidermis. We utilize single-cell ribosome profiling to quantify mRNA translation in the different cell types of the epidermis, annotate cell-type-specific noncanonical open reading frames and codon occupancies. We describe the heterogeneity in the epidermal stem cell population and the switches in translational efficiency during differentiation, unveiling basic posttranscriptional processes in epidermal biology. By coupling single-cell ribosome profiling with single-cell RNA sequencing, we then, for the first time, document the cell-type-specific *in vivo* changes of the translational efficiency landscape in aging. We uncover translational reprogramming of AP-1 subunits in aging specifically in epidermal stem cells and suprabasal cells. Finally, we provide functional evidence that AP-1 overexpression induces aging-relevant changes in keratinocyte function.

AP-1 levels control inflammatory responses^50^ and inflammaging^51^. Higher AP-1 protein levels in aging also play a critical role in chromatin opening, which disrupts the binding of cell identity transcription factors to cis-regulatory elements and can siphon cell identity transcription factors away from their typical regulatory elements, thereby altering cellular function^52^. Thus, our findings prompt further studies into the posttranscriptional regulation of AP-1 in aged stem cells, examining how aging-related changes in the translational machinery elevate AP-1 and how AP-1 impacts *in vivo* stem cell function and inflammatory responses in aging. In contrast, several genes critical for epidermal biology, including E-cadherin, semaphorins and calreticulin exhibit a marked downregulation in translational efficiency, impacting tissue-wide cell-cell signaling (Figure 4Q-S) and potentially contributing to tissue function decline during aging. These findings underscore the power of *in vivo* single-cell ribosome profiling data in generating tissue-level hypotheses, advancing our understanding of posttranscriptional regulatory processes in aging and uncovering novel entry points for therapies to counteract the decline of stem cell function.

Collectively, our strategy illustrates the utility of *in vivo* single-cell ribosome profiling to document cell-type-specific translational landscapes, reveals widespread translational reprogramming in aged stem cell populations and provides the basis into whether therapies interfering with the aging translatome could preserve stem cell function in aging.

## Acknowledgements

We thank Ruben Barricarte, Felipe G. Quiroz, and Sendoel lab members for critical input on the manuscript, Catharine Aquino and the FGCZ for sequencing, the LASC Zurich for assistance with mouse work and Philipp Schätzle and the UZH Center for Microscopy and Image Analysis for assistance with microscopy. The project was supported by the European

Research Council (ERC) under the European Union′s Horizon 2020 research and innovation programme (grant agreement No 759006), by the Helmut Horten Foundation, by an SNSF Professorship grant (grant number 176825) and the National Center of Competence in Research (NCCR) on RNA and Disease funded by the SNSF (grant number 205601).

## Author contributions

C.D. conducted the experiments and collected the data. U.G. performed bioinformatic data processing. C.D., R.W and A.S. developed the single-cell ribosome profiling protocol. C.D., U.G., R.W., D.T. and A.S. performed data analysis and interpretation. F.V.-F. assisted with mouse preparations, cloning and lentiviral preparations. P.F.R. sorted cells for single-cell ribosome profiling. A.K. and S.J.E. performed immunofluorescence stainings and image analysis. F.S. and M.L. performed immunohistochemistry stainings. K.H. assisted with cloning. M.Y. provided protein samples. M.O. and H.Y. assisted with animal experiments.

C.D. and A.S. conceived the project. A.S. supervised the project.

## Material & Methods

### *In vivo* experiments

All animal experiments were conducted in strict accordance with the Swiss Animal Protection law and requirements of the Swiss Federal Office of Food Safety and Animal Welfare (BLV). The Animal Welfare Committee of the Canton of Zurich approved all animal protocols and experiments performed in this study (animal permits ZH233/2019 and ZH152/2023).

### Mouse preparation for ribosome profiling

Cycloheximide (CHX; 20 mg/ml, Sigma-Aldrich) was injected into the tail vein of female young (2 month) and old (22-26 month) mice prior to euthanasia, as previously described.^19^ The back skin was surgically removed and submerged in 1x PBS (Gibco; 10010-015), supplemented with CHX (10 mg/ml) for 5 minutes. The fat was scraped from the back skin and washed once in cold PBS-CHX (0.1 mg/ml CHX) and then placed in 15 ml 0.25% trypsin (2 mg/ml CHX) (Thermo Fisher Scientific, 25050014) for 37 minutes at 37 °C on an orbital shaker, dermis-side facing down. Next, the epidermis was separated from dermis by scraping with a scalpel and torn into smaller pieces. Trypsin was neutralized by adding 1x PBS buffer containing 2% chelexed FBS (FBS-) and CHX (0.1 mg/ml), followed by vigorous pipetting to create a single-cell suspension. The cell suspension was filtered sequentially through a 70 µm and a 40 µm strainer (Corning; 431750, 431751), centrifuged for 10 minutes at 400 g and then resuspended in FACS buffer (1x PBS, 2% FBS-, 0.1 mg/ml CHX, 0.05 ng/ml DAPI).

### Cell lysis and footprint generation

Live single cells were sorted on a BD Aria II with a 70 µm nozzle and a flow rate of <300 events per second into 384-well hard-shell plates (Bio-Rad, HSP3805), pre-filled with 5 µl of light mineral oil (high purity, Alfa Aesar) and 50 nl of lysis buffer (22 mM Tris-HCl pH 7.5, 16.5 mM MgCl2, 165 mM NaCl, 1.1% Triton X-100, 0.1 mg/ml CHX). After sorting, plates were spun down at 4,000 g for 2 minutes and maintained on wet ice until all plates were prepared for further processing. Next, 50 nl of RNase I (0.2 U/µl lysate; 500 U/µg RNA, New England Biolabs) was added to each well, and plates were incubated at room temperature for 45 minutes. To stop digestion, 50 nl of stop mix (0.0186 U/µl Thermolabile Proteinase K (New England Biolabs), 62 mM EGTA (Sigma-Aldrich), 16.5 mM EDTA (Ambion), 697.5 mM guanidium thiocyanate (GuSCN, Sigma-Aldrich)) and 0.8 U/µl SUPERase·In™ RNase Inhibitor (Invitrogen, AM2696) were added to each well, and plates were incubated at 37 °C for 30 minutes, followed by 55 °C for 10 minutes and held at 4 °C.

### Nuclease testing

RNase T1 (Thermo Fisher Scientific, EN0542) and RNase I (Biosearch Technologies, N6901K) were tested in different concentrations to evaluate the ideal concentration to generate single-cell ribosome profiling libraries. RNase T1 digestion was performed using 10, 150 and 450 U/µg RNA. RNase I digestion was performed using 0.5, 5, 50 and 500 U/µg RNA, with the 500 U/µg RNA concentration defined as the optimal concentration for nuclease footprinting in single-cell ribosome profiling.

### Ribosome profiling library preparation

After ribosome footprint generation, libraries were generated using the Mosquito® liquid handler (SPT Labtech). As previously described^15^, a one-pot small-RNA library preparation protocol included the end repair, two RNA ligations, cDNA synthesis, and an indexing PCR.

End repair: First, 50 nl of end-repair mix (4.1× T4 RNA Ligase Buffer (New England Biolabs), 16.4 mM MgCl2, 4.1 mM uridine triphosphate (New England Biolabs), 1.37 U/µl T4 Polynucleotide Kinase (New England Biolabs) and 0.77 U/µl SUPERase·In™ RNase Inhibitor was added to each well, and plates were incubated at 37 °C for 1 hour, then held at 4 °C.

3’ Ligation: Next, 264 nl of 3’ ligation brew (1× T4 RNA Ligase Buffer (New England Biolabs), 1 µM pre-adenylated 3’ linker (Integrated DNA Technologies), 35.5% PEG-8000 (New England Biolabs), 0.1% Tween-20 (Sigma-Aldrich), 0.98 U/µl SUPERase In RNase Inhibitor (Invitrogen), and 21.3 U/µl T4 RNA Ligase 2 Truncated KQ (New England Biolabs) was added to each well and plates were incubated at 4 °C for 18 hours.

To prepare 20 µM pre-adenylated linker, 1.2 µl of linker oligonucleotide (Integrated DNA Technologies) were mixed with 2 µl 10x 5’DNA adenylation buffer (New England Biolabs, E2610L), 0.095 mM ATP, 13.8 µl H2O and 20.56 µM Mth RNA ligase (New England Biolabs, M2611A). The mix was incubated at 65°C for 60 minutes followed by an inactivation step at 85°C for 5 minutes. The pre-adenylated linker was eluted in 6 µl H_2_O following the clean-up using the Oligo Clean & Concentrator Kit (Zymo Research, D4060).

The cDNA synthesis primer was then pre-annealed to the 3’ ligation products by adding 50 nl of the reverse transcription primer mix (5.2 µM reverse transcription primer (Integrated DNA Technologies), 13.5 µM adenosine triphosphate (ATP, New England Biolabs), and 1% Tween-20 to each well. The plates were heated to 65 °C for 1 minute, then incubated sequentially at 37 °C for 2 minutes, 25 °C for 2 minutes and finally held at 4 °C.

5’ Ligation: 5’ adapters were then ligated by adding 156 nl of 5’ ligation brew (1× T4 RNA Ligase Buffer, 30.75% PEG-8000, 0.1% Tween-20, 0.5 µM 5’ adapter (Integrated DNA Technologies), 1.25 U/µl T4 RNA Ligase (Thermo Scientific) and incubated at 37 °C for 2 h and then held at 4 °C.

cDNA synthesis: Complementary DNA synthesis was performed by adding 770 nl of reverse transcription reaction with 2.66x of the standard concentration of the RT Buffer (5x RT Buffer, Thermo Fisher Scientific, EP0752), 1.25 mM dNTPs (Promega), 0.19% Tween-20, 1.88 U/µl SUPERase In RNase Inhibitor, and 9.4 U/µl Maxima H Minus Reverse Transcriptase (Thermo Fisher Scientific) to each well. The plate was incubated at 50 °C for 1 hour, followed by 85 °C for 5 minutes, then held at 4 °C.

Indexing PCR: For library indexing, 150 nl of 20 µM unique forward index primers (Integrated DNA Technologies) and 3.2 µl of PCR mix (1.5× Q5 Hot Start High-Fidelity 2× Master Mix (New England Biolabs), 0.15% Tween-20, and 0.94 µM reverse index primer (Integrated DNA Technologies)) were added to each well. Plates were then incubated at 98 °C for 30 seconds followed by 10 cycles of 98 °C for 15 seconds, 65 °C for 30 sesonds, 72 °C for 30 seconds, and then a final incubation at 72 °C for 5 minutes. Plates were then held at 4 °C before freezing at −20 °C until pooling.

### Pooling and purification

Folllowing library construction, the plates were pooled and purified. The contents of each plate were first collected in an 86 × 130 mm container (1000 µl pipette tip lid) by centrifuging at 800 g for 2 minutes and transferring into a 15 ml tube. The aqueous phase (∼1.9 ml per plate) was separated from the light mineral oil by centrifugation at 2,000 g, and then concentrated to approximately 500 µl using n-butanol (Sigma-Aldrich). The product was then cleaned up using AMPure XP beads (Beckman Coulter), pre-diluted 5x in bead binding buffer (20% PEG-8000 (Sigma-Aldrich) 2.5 M sodium chloride (Sigma-Aldrich)). Diluted beads were added to the sample at a 2.1:1 ratio, and the final product was resuspended in 50 µl low TE buffer (LoTET, 3 mM Tris-HCl pH 8.0 (Ambion), 0.2 mM EDTA pH 8.0 (Gibco), 0.1% Tween-20). 10 µl of each of the cleaned library pools was then run on a 17.5-cm 7% polyacrylamide gel at 120 V for ∼6 h. The ∼10-bp region corresponding to an insert size of ∼30–40 nt was excised from 175 bp to 185 bp of the polyacrylamide gel. The excised band was then crushed and resuspended in 400 µl DNA gel extraction buffer (10 mM Tris-HCl pH=8, 300 mM NaCl, 1 mM EDTA), frozen at -80°C for 30 minutes and eluted overnight at room temperature with gently shaking. The liquid was recovered into a new tube. To precipitate the DNA, 1.5 µl GlycoBlue coprecipitant (Invitrogen) and 500 µl isopropanol were added to the extracted DNA solution, followed by overnight incubation at −80 °C. The sample was centrifuged at 20,000 g for 1 hour at 4 °C. The supernatant was carefully removed, and the pellet was air-dried before being resuspended in 15 µl of 10 mM Tris-HCl, pH 8.0.

### Sequencing

Libraries were sequenced on a NovaSeq 6000 (Illumina) or NovaSeq X Plus (Illumina), using a minimum of 75 cycles for read 1, 6 cycles for the i7 index read (plate index), and 10 cycles for the i5 index read (cell index).

### Single-cell RNA sequencing

Epidermal cells were isolated from one young (2 month old) and one aged (24 month old) animal as described earlier and processed for single-cell capture with a BD Rhapsody Single-Cell Analysis System with the BD Rhapsody Cartridge Kit^45^ (633733). The cell suspension was barcoded using BD Single-Cell Multiplexing Set (Rat anti-mouse MHC-H2 Class I, BD Biosciences, 626545), and 40,958 cells were loaded into the cartridge. Reverse transcription from beads and sequencing library production was carried out according to the manufacturer’s instructions (BD Biosciences, doc ID: 210967 rev. 1.0). After processing through the BD bioinformatics pipeline, 15,873 cells were retrieved (young = 7,426 cells, old = 8,447 cells). RNA single-cell sequencing was performed on an Illumina NovaSeq 6000 SP with the following sequencing configurations: Read 1 60 cycles, index 1 8 cycles, read 2 62 cycles. 1.8nM of the library was loaded with 20% PhiX.

### Histology and immunofluorescence

Fixed back skin pieces cut as strips along the AP axis were embedded in OCT (Tissue-Tek; 4583) and frozen over dry ice. 10 µm thick sagittal sections were obtained using a Micron HM500 OM cryostat (Thermo Scientific) and immobilized on Superfrost glass slides. Slides were washed twice in 1× PBS before blocking in blocking solution (1% BSA, 1% gelatin, 2.5% normal goat serum, 2.5% normal donkey serum, 0.30% Triton X-100, 1× PBS). Primary antibodies CD324 (E-Cadherin) anti-rat antibody (Invitrogen, 14-3249-82), 1:200, c-Jun anti-rabbit antibody (60A8, CST 9165), 1:50, and anti-mouse Keratin 6A Antibody (Biolegend 905702), 1:200 were incubated overnight at 4 °C. After washing twice with 1× PBS, slides were incubated with secondary antibodies (Alexa Fluor 488, Cy3, or Alexa Fluor 647, Jackson ImmunoResearch Laboratory; 1:500) and 1 µg/ml 4′,6-diamidino-2-phenylindole (DAPI) at room temperature for 1 hour. Sections were then washed again with 1× PBS, dried, covered with ProLong™ Diamond Anti-fade Mountant (Invitrogen, P36965) and sealed with a coverslip and allowed to cure for 24 hours before imaging.

### Imaging and image processing

Tiled images were acquired on a Zeiss LSM900 microscope using a LD LCI Plan-Apochromat 40x/1.2 Multi-Immersion, WD 0.41 mm objective with glycerine as an immersion medium. Images were processed (stitched) using Zen Blue. Further processing and quantifications (Jun intensities) were done in Fiji (ImageJ). All images depicted are maximum intensity projections.

### Quantification of nuclear JUN intensity

The DAPI channel in the maximum intensity projections was used to create nuclear masks. Briefly, thresholded nuclei were detected using the Analyse particles tool in Fiji. Accuracy of detection was checked manually and adjusted for (by removing or adding particles) as needed. The generated masks were used to measure mean grey values in the Jun channel. Nuclei from the dermal compartment and hair follicles were excluded from analysis. An unpaired two tailed t test was performed and data were plotted using GraphPad Version 9.5.0 for Mac OS, GraphPad Software, Boston, Massachusetts USA.

### Protein isolation for western blotting

Epidermal cells were isolated from young (2 month old) and aged (26 month old) animals as described above. The cell suspension was stained with Ly-6A/E (Sca-1) Monoclonal Antibody, FITC (1:100, Thermo Fisher Scientific, 11-5981-82) for 30 minutes at 4°C, followed by a wash with FACS buffer and resuspension in FACS buffer supplemented with DAPI (0.05 ng/ml).

Cells were sorted on a BD FACSAria™ III Cell Sorter using a 70 µm nozzle. Cells were collected in E-low medium and centrifuged at 400 g for 15 minutes. The cell pellet was lyzed in RIPA buffer (Sigma-Aldrich, R0278), supplemented with Protease Inhibitor Cocktail (50x, Promega, G6521), Phosphatase inhibitor cocktail 2 (100x, Sigma, P5726-1ML) and Phosphatase inhibitor cocktail 3 (200x, Sigma, P0044-1ML) and incubated on ice for 10 minutes. The cell lysate was centrifuged at full speed for 10 minutes at 4°C and supernatant was collected. Protein concentration was measured using the Pierce BCA Protein Assay Kit (Thermo Fisher Scientific, 23225). Protein lysates were mixed with 4x sample loading buffer (0.2 M Tris-HCL, 0.4 M DTT, 8% SDS (w/v), 6 mM bromphenol blue, 4.3 M glycerol) and heated to 95°C for 10 minutes, then stored at -80°C.

Protein samples were re-heated to 95°C for 10 minutes and loaded on a 4-12% NuPage 12-well Bis-Tris Gel (Invitrogen, 7001691) using NuPAGE MOPS SDS Running Buffer (20X, Invitrogen, NP0001). Proteins were subsequently transferred to a nitrocellulose membrane (GE Healthcare) by tank transfer at 4°C, 30V for 1.5 hours. Membranes were blocked in 5% skimmed milk. Primary antibodies were incubated overnight at 4°C and secondary antibodies for 2 hours at 4 °C. Western blots were developed with freshly mixed ECL solutions (GE Healthcare). The following antibodies were used: c-Jun (60A8) Rabbit mAb (Cell Signaling Technology, 9165), Phospho-4E-BP1 (Thr37/46) (Cell Signaling, 2855), Anti-Vinculin (Abcam, ab129002) from Sigma Aldrich (T6199). Western blot bands were quantified using ImageJ.

### CROP-mCherry vector and cloning

The ampicillin resistance cassette in the original CROP-seq-Guide-Puro vector (Addgene #86708) was replaced with an mCherry sequence, amplified from pAAVS1-NDi-CRISPRi (Gen1) (Addgene #73497) and cloned in via Pfl23II and MluI restriction sites. The Crop-mCherry plasmid was digested with BsmBI to remove the filler sequence and replaced with a nontargeting sgRNA. *Fos* gBlock (Ref #510140372) was obtained from IDT and the restriction sites BamHI and MluI, along with a 2A site were added via PCR using IDT primers. The plasmids were sequenced to confirm correct insertion of *Fos*.

### Primer sequences

Fos fw: ATGGATCCGGGgccaccaacttcagcctgctgaagcaggccggcgacgtggaggagaaccccggccccATGATGTTCTCGGGTT Fos rev: taacgcgtTCACAGGGCCAGCAGCGTG

### Primary keratinocyte isolation and culture

Primary mouse epidermal keratinocytes were isolated from 4 day old C57BL/6 wildtype mice, as previously described.^53^ Briefly, isolated epidermal keratinocytes were cultured on 3T3-S2 feeder layer pre-treated with Mitomycin-C, using 0.05 mM Ca^2+^ E-media supplemented with 15% serum. After 3 passages on 3T3-S2 feeder layer, cells were transferred to 0.05 mM Ca^2+^ E-media, prepared in-house, as previously described^54^. The C57BL/6 wildtype keratinocytes were used for protein sample collection, lentiviral infection and scratch assays. All cell cultures were maintained under standard conditions at 37 °C and 5% CO_2_.

### Low-titer lentivirus production

Production of vesicular stomatitis virus (VSV-G) pseudotyped lentivirus was performed by calcium phosphate transfection of Lenti-X™ 293T cells (TaKaRa Clontech, 632180) using the CROP-mCherry-Fos construct and the helper plasmids pMD2.G and psPAX2 (Addgene plasmids 12259 and 12260), as previously described. Viral supernatant was collected 46 h after transfection and filtered through a 0.45 µm filter (Sarstedt AG, 83.1826).

### High-titer lentivirus production

High-titer lentivirus production followed the same procedure as for low-titer lentivirus, with an additional concentration step. The viral supernatant was concentrated ∼2,000-fold using a 100 kDa MW cut-off Millipore Centricon 70 Plus (Merck Millipore, UFC710008). The virus was further concentrated by ultracentrifugation using the SW 55 Ti rotor for the Beckman Coulter Optima L-90 Ultracentrifuge at 45,000 rpm at 4 °C. Viral particles were resuspended in viral resuspension buffer (20 mM Tris pH 8.0, 250 mM NaCl, 10 mM MgCl2, 5% sorbitol) and stored at −80 °C until used for infection.

### Transfection

Keratinocytes were transfected with CROP-mCherry-Fos using lipofectamine 2000 according to manufacturer’s recommendation. Briefly, 100 ng of DNA was transfected using Opti-MEM and Lipofectamine 2000 Reagent.

### *In vitro* lentivirus infections

For lentiviral infections in culture, C57BL /6 wildtype keratinocytes were seeded in a 6-well plate (Thermo Scientific Nunclon TM Delta Surface; 140675) at 1.5-3 × 10^5^ cells per well. Cells were infected with 100-300 µl of low-titer virus in the presence of the infection mix (1/10 dilution of polybrene [10 mg/ml Sigma; 107689-100MG in PBS] in FBS[-]), followed by centrifugation of the plates at 1,100 g for 30 minutes at 37 °C in a Thermo Heraeus Megafuge 40R centrifuge. Infected cells were FAC-sorted for mCherry on a BD Aria II.

### Scratch assay

Cells were seeded in an ibidi Culture-Insert 2-well (81176) according to the manufacturer’s recommendations. Lentivirus-infected or transfected keratinocytes overexpressing *Fos* and control keratinocytes were used for the scratch assay. The culture insert creates a clean cell-free gap in a 24-well plate (VWR, 734-2325). The assay was performed according to the manufacturer’s protocol (ibidi). Briefly, once cells reached confluency, the insert was removed, and detached cells were washed away with 1x PBS. New medium was added, and the wells were imaged every hour for 24 hours in an incubation chamber at 37 °C and 5% CO2. The images were taken with a Zeiss Axio Observer controlled by the ZEN software. Data analysis was carried out with an ImageJ plugin67 (version 1.54h) using default settings.

### Immunofluorescence of protein synthesis

For protein synthesis analysis, O-Propargyl-puromycin (OPP, Jena Biosciences, NU-931-5, 50 µg/g body mass, in PBS) was administered via intraperitoneal injection into P4 animals 1 hour before euthanasia. Skin tissues were then embedded, sectioned, and processed for staining as described in the immunofluorescence section, followed by staining with Alexa Fluor 594 azide (1:500, Life Technologies). Image quantification (OPP intensities) was done in Fiji (ImageJ).

### Demultiplexing of the single-cell ribosome profiling data

Raw Illumina sequencing data were demultiplexed using the bcl2fastq software (version 2.20.0.422) with the options --barcode-mismatches 1 -- minimum-trimmed-read-length 5 --mask-short-adapter-reads 5. The demultiplexing process used a CSV file where the cell barcode was indexed under “index2” and the sample barcode under “index.” This configuration ensured that the resulting FASTQ files contained both sample and cell barcodes in the header for downstream processing.

### Processing of single-cell ribosome profiling data

We used fastp to trim adapters and relocate the 10nt UMI sequence to the FASTQ header, retaining reads with lengths between 26-34 nucleotides. Next, rRNA and tRNA sequences were removed using bowtie2. A custom Python script was then employed to generate a FASTQ file that included the 10nt cell barcode and 10nt UMI sequence. The resulting fastq files were aligned to the mouse genome using STAR. The resulting bam file was cleaned up for empty cell barcodes with sambamba followed by UMI deduplication by umi_tools. The deduplicated bam files were split into 384 cells by a custom python script based on the pysam library. FeatureCounts was used to quantify reads mapped to coding sequences (CDS) regions.

### Quality control of the single-cell ribosome profiling data

Quality control was conducted using the riboWaltz R package^56^. The deduplicated transcriptome BAM files were converted using the bamtolist command and M25 gencode annotations for transcripts with CDS length > 0. The p-site offsets were computed via the psite command in automatic mode with flanking 6. Periodicity and other QC plots were generated with riboWaltz commands.

### Detection of noncanonical open reading frames (ORFs)

We utilized the Ribotricer software to detect novel ORFs from the single-cell ribosome profiling data^26^. First, the prepare-orfs subcommand was used with the GRCm38 primary assembly genome reference and gencode M25 annotations, focusing on canonical ATG start codons and a minimum ORF length of 30 nucleotides (10 amino acids). The Ribotricer “detect-orfs” command was then run for each cluster of young and old skin cells, specifying read lengths between 26-34 nucleotides, forward strand mode, and a p-site offset of 12.

### CellChat

To analyze cell-to-cell signaling between clusters, we utilized the CellChat package^42^ with the truncated mean approach, requiring a minimum of 10 cells for inferring signaling interactions.

### Single-cell RNA data processing

The raw sequencing data, provided as BCL files, were demultiplexed with Illumina’s bcl2fastq software (v2.20.0.422) using default settings, allowing for a single mismatch in the sample barcode. The resulting FASTQ files were then processed using the BD Rhapsody Whole Transcriptome Analysis (WTA) Pipeline (v1.11) on the Seven Bridges Genomics cloud platform. A STAR (v2.5.4) reference genome was created with Gencode mouse vM25 (GRCm38.p6) annotations for use in the BD pipeline. Young and old skin samples were multiplexed with the BD Single-Cell Multiplex Kit for Mouse, using barcodes 5 and 6, respectively. The BD pipeline’s Multiplexing_Settings option was configured for the Single-Cell Multiplex Kit - Mouse to facilitate sample demultiplexing. UMI counts, corrected using BD’s Recursive Substitution Error Correction (RSEC) method, were applied in subsequent analyses.

### Analysis of the single-cell RNA sequencing data

First, Scrublet library from the Scanpy Python package was used to identify doublets in the young and old skin scRNA-seq data. Next, we removed the cells identified as doublets and merged the samples in Seurat using the merge command. We selected cells with counts >500 and <6000 and retained only cells with <20% mitochondrial gene content. Next, we employed the standard Seurat pipeline to process the data. First, the counts were normalized by the NormalizeData function to scale the counts to 10,000 per cell followed by ln(counts+1). We ran FindVariableGenes function to detect the 2000 most variable genes. The normalized data was scaled by the ScaleData function. Next, dimensionality reduction by Principal Component Analysis (PCA) was performed with the RunPCA function. We analyzed the contribution of each Principal Component for explaining the proportion of variance in the data with the ElbowPlot function and selected the top 50 principal components. Next, nearest neighbors were identified by the FindNeighbors function. We carried out the clustering of the data with FindClusters function with resolution 0.2 using modularity optimization by Louvain algorithm. Uniform Manifold Approximation and Projection (UMAP) dimensional reduction technique was used to visualize the data. The young and old samples were clustered together in the UMAP showing no batch effect as the samples were prepared together. We obtained the marker genes for each cluster by the FindAllMarkers function and annotated the clusters with cell types based on the expression of known marker genes. We used Nebulosa to visualize the gene expression data with kernel density estimation.

### Differential expression analysis

Single-cell differential expression (DE) analysis was conducted using the glmGamPoi R package, which fits Gamma-Poisson models to the data. First, we selected genes expressed in at least 10% of cells, following a similar threshold as in the Seurat DE approach. We then fit a Gamma-Poisson generalized linear model to the data using the glm_gp function with default settings, allowing both overdispersion and overdispersion shrinkage. A quasi-likelihood ratio test for the Gamma-Poisson fit was performed using the test_de function to obtain p values for differential expression.

Pseudobulk DE and direct TE analysis were carried out by DESeq2. First, the normalization factors for each sample were computed by the estimateSizeFactors function using the median of ratios method. Next, the estimation of dispersion was performed by the estimateDispersions function. A negative binomial generalized linear model was fitted for each gene to identify differentially expressed (DE) genes. The p values were obtained by the Wald test and were corrected for false discovery rate using the Benjamini-Hochberg procedure.

### Translational efficiency (TE) analysis

TE analysis between old vs young skin across the clusters were carried out using a pseudobulk approach with DESeq2. Each cluster from young and old skin scRNA-seq and scRibo-seq data was randomly divided into 3 groups and raw UMI counts were aggregated to generate pseudobulk data. First, the normalization and dispersion estimation were performed as in the DE analysis. Next, TE for old vs young skin for each cluster was computed using the negative binomial likelihood ratio test by the nbinomLRT function with ∼ assay + condition + assay:condition as full model and ∼ assay + condition as reduced model. Assay refers to single-cell RNA sequencing and single-cell ribosome profiling data, while condition refers to old and young epidermis. Genes with mean counts >10 were selected for TE analysis.

### TE and TE permutation test

Cell-type-specific TE was computed as log_2_((scRibo-seq_TPM + 1) / (scRNA-seq_CPM + 1). We performed 10,000 permutations to calculate p values for translational efficiency (TE) changes per cluster in young skin using single-cell ribosome profiling and single-cell RNA sequencing data. First, UMI counts from both datasets were filtered to retain only protein-coding genes. Pseudobulk values were then computed, with single-cell ribosome profiling data expressed as transcripts per kilobase million (TPM) and single-cell RNA sequencing data as counts per million (CPM). TE was defined as log_2_((scRibo-seq_TPM + 1) / (scRNA-seq_CPM + 1)). Next, we randomly sampled an equal number of cells from single-cell ribosome profiling and single-cell RNA sequencing in the merged data set 10,000 times to generate a pseudobulk TE null distribution. Two-sided empirical p-values for each gene were calculated by determining how often the absolute value of the null distribution TE (|TEnull|) exceeded or equaled the absolute observed TE (|TE|), with the formula: p = (|TEnull| ≥|TE|) / 10,000.

### General data analysis

The analysis was carried out using in-house Python 3.9 and R 4.1 scripts. Data wrangling was done with the Pandas library in Python and tidyverse library in R. Heatmaps were constructed using the ComplexHeatmap and pheatmap packages. Other plots were made using the ggplot2 library in R and seaborn library in Python.

### Pathway enrichment

Pathway enrichment analysis was performed using EnrichR^57^ (https://maayanlab.cloud/Enrichr), which applies Fisher’s exact test, with custom R scripts leveraging the enrichR library for GO terms and other gene sets. A cut-off of FDR < 0.05 was used to select differentially expressed genes.

Additionally, we employed the pre-ranked method of GSEA^58^ and ssGSEA with GSEApy^59^ (https://github.com/zqfang/GSEApy) to assess both up- and down-regulated genes simultaneously, enhancing the sensitivity of gene set enrichment analysis. We conducted 1,000 permutations and used an FDR cutoff of 0.05 to determine enriched gene sets.

**Figure S1:**
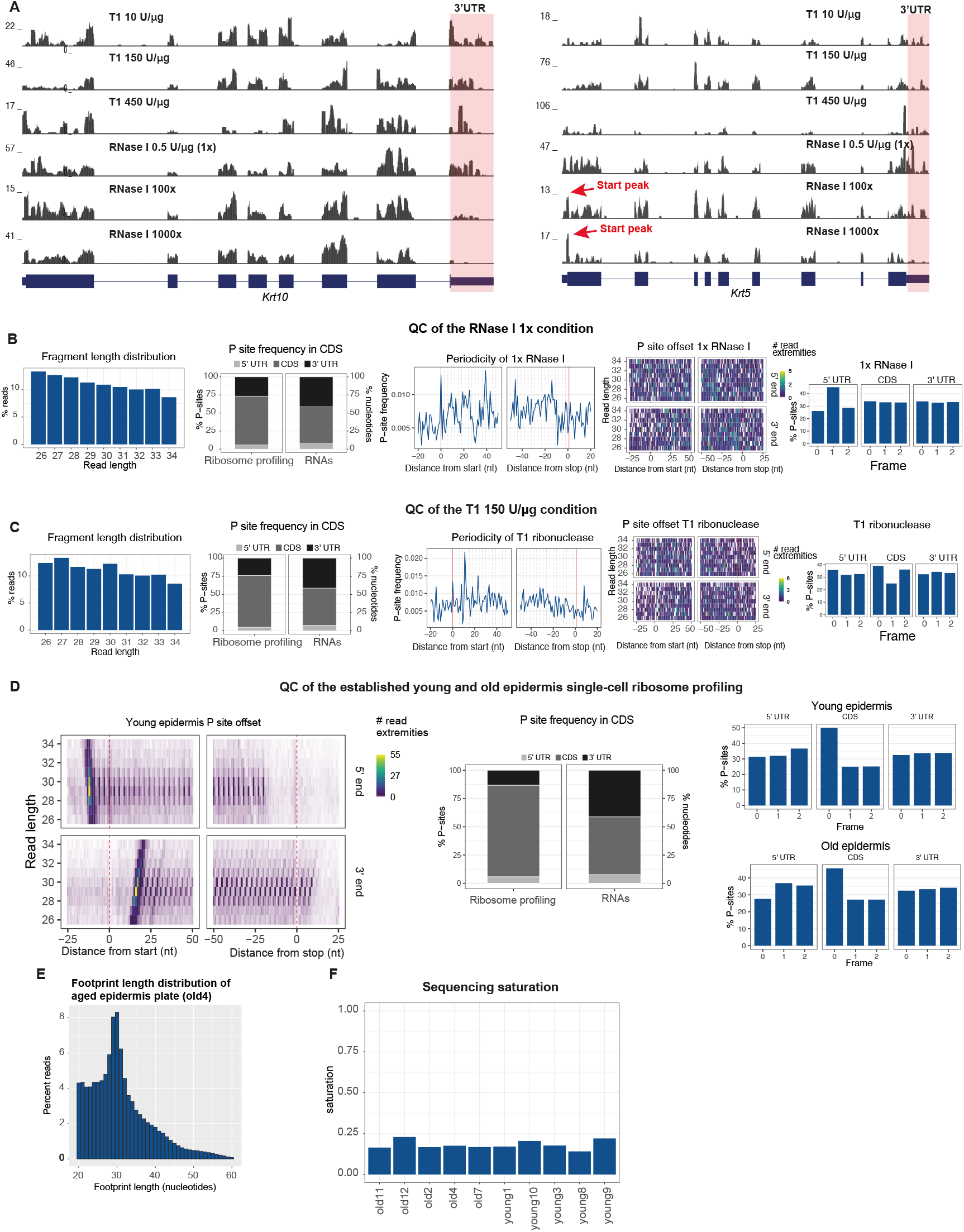
Quality control of the *in vivo* single-cell ribosome profiling protocol. (A) Ribosome profiling tracks *of Krt10* or *Krt5* following different concentrations of the ribonucleases RNase T1 and RNase I reveal stepwise reductions of the 3’UTR reads, with the cleanest pattern in the 1000x RNase I condition. (B-C) Quality control (QC) of single-cell ribosome profiling tests using either 1x RNase I (B) or RNase T1 (150U/μg) (C) concentrations, analyzed with riboWaltz, reveals suboptimal fragment length distributions, a high proportion of reads mapping to the 3’UTR, lack of 3-nucleotide periodicity or clear start site peak, noisy P-site offset and no CDS frame signals (from left to right panels). (D) Quality control of established single-cell ribosome profiling with 1000x RNase I concentration shows a clear P site offset signal, a start site peak, the majority of reads mapping to the CDS and a clear CDS frame. (E) Footprint length distribution from single-cell ribosome profiling of aged epidermis reveals a peak at 29 nucleotides, similar to that observed in young animals. (F) The sequencing saturation of the main plates processed suggests that the single-cell ribosome profiling approach is scalable, with the potential to increase unique read count distribution and number of quantified genes through additional sequencing. Saturation is defined as: 1 - (Number of unique reads/ Total number of reads).

**Figure S2:**
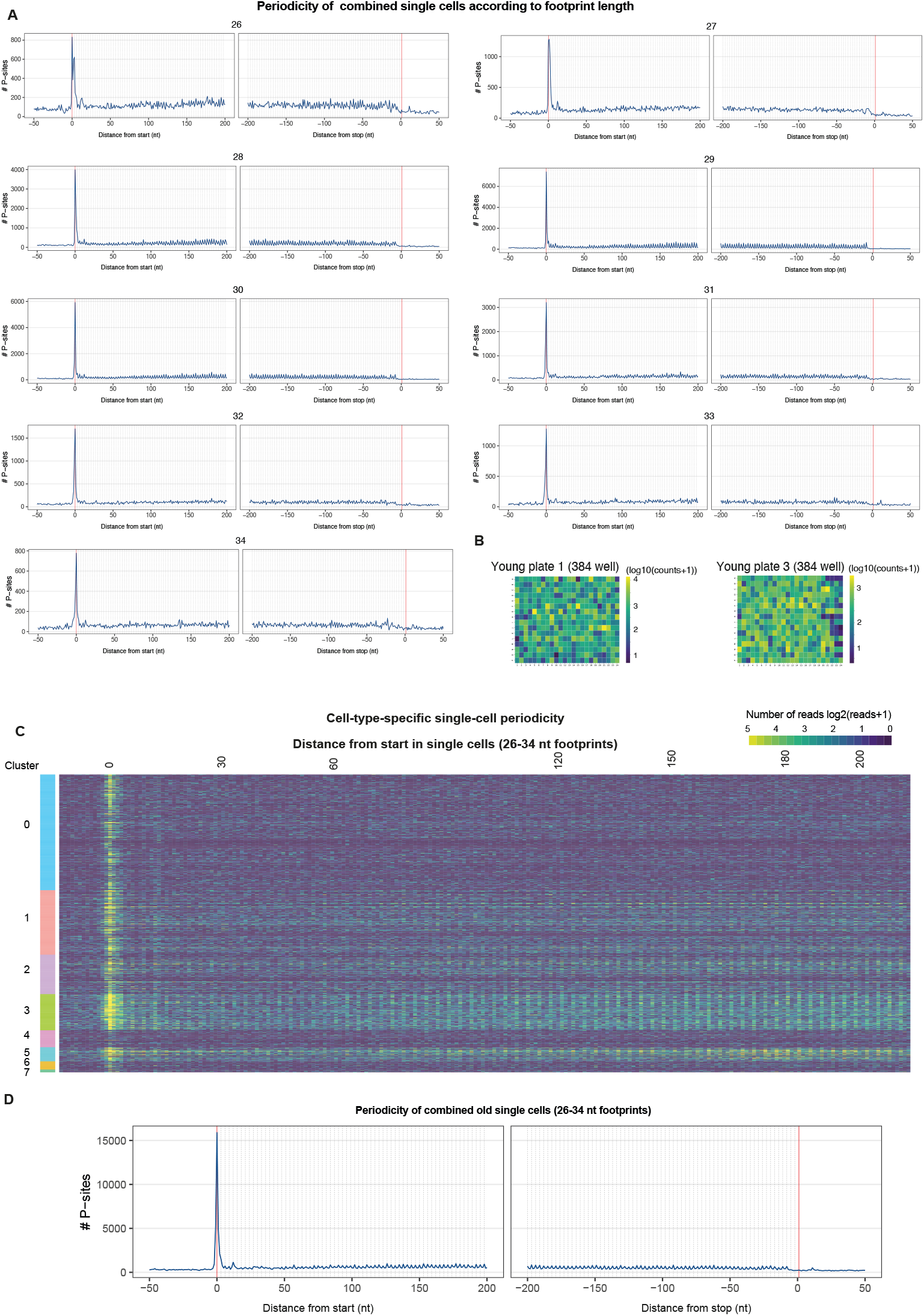
Detection of triplet periodicity in the *in vivo* single-cell ribosome profiling data. (A) Triplet nucleotide periodicity analysis of pooled single cells shows the most distinct periodicity for the footprints between 28 and 31 nucleotides in length. P sites are mapped across the coding sequence 200 nucleotides downstream of the start codon and 200 nucleotides upstream of the stop codon. (B) Heatmap of UMI counts of two representative plates filled with young (2 months) epidermal cells show overall uniform sequencing depth across the plate. (C) Heatmap of cell-type-specific triplet nucleotide periodicity (mapped P sites) of single cells highlights low overall UMI counts in epidermal stem cells (cluster 0) and bulge stem cells (cluster 4). Clusters refer to Figure 2A. (D) Triplet periodicity of combined old single cells with 26-34 nucleotide fragment length demonstrates robust periodicity across the coding sequence.

**Figure S3:**
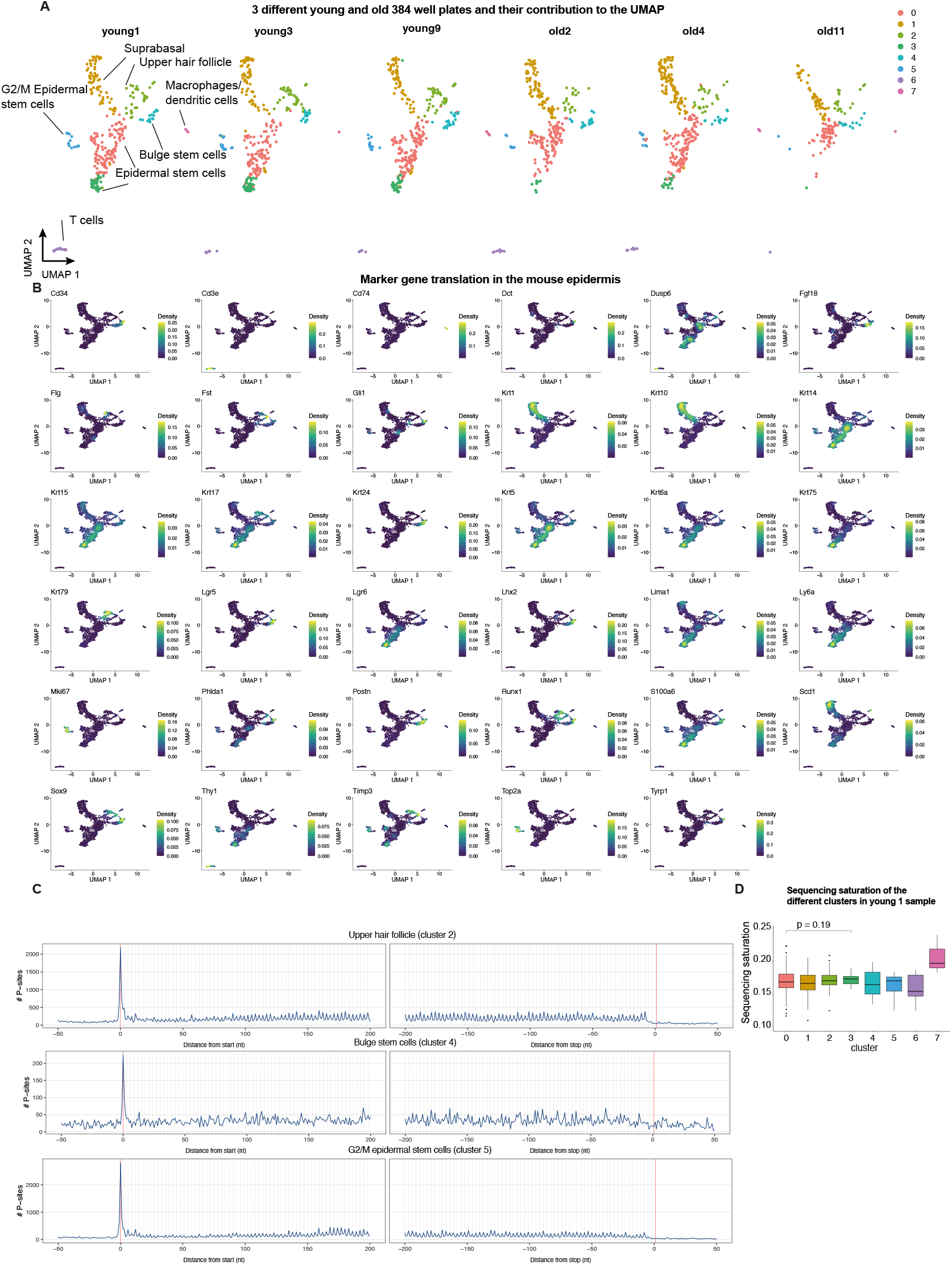
Marker gene translation across different epidermal cell types. (A) The contribution of cells from three 384-well plates from young mice and from three 384-well plates from old mice to the UMAP demonstrates an overall balanced representation of cell types across the young and old plates. (B) Nebulosa density plots of marker gene translation in the mouse epidermis. (C) Triplet periodicity of different cell types, including the upper hair follicle, bulge stem cells and G2/M epidermal stem cells of the young mouse skin. (D) The sequencing saturation levels for the cells of clusters 3 and 5 (high number of unique reads) do not exceed that of cluster 0. Saturation is defined as: 1 - (Number of unique reads / Total number of reads) calculated for the young 1 sample. P value indicates a Wilcoxon test.

**Figure S4:**
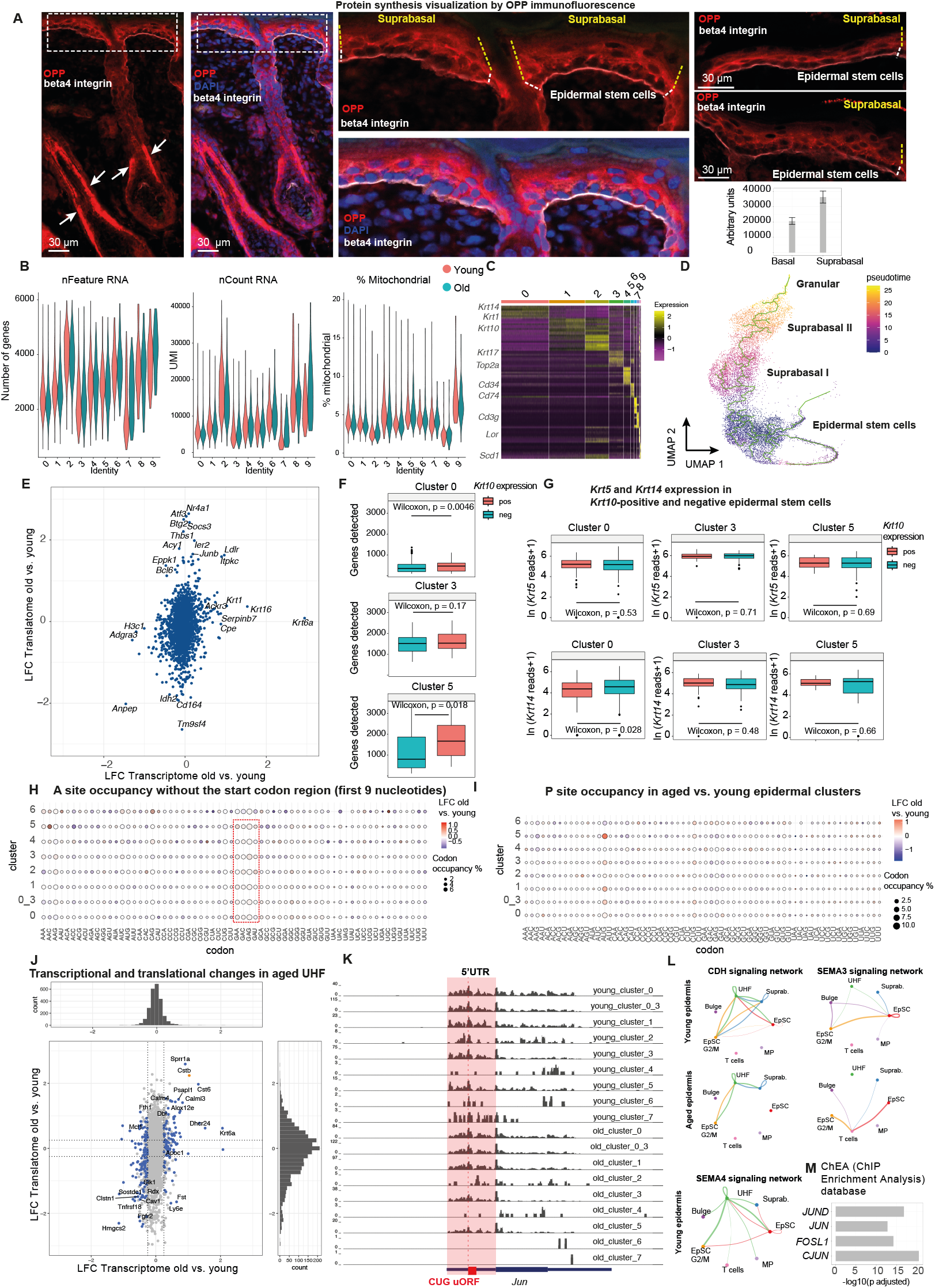
The translational landscape of the epidermis. (A) O-propargyl-puromycin (OPP) incorporation and immunofluorescence of the mouse skin exposes lower protein synthesis rates in epidermal stem cells (white dashed line) compared to suprabasal cells (yellow dashed line). P4 animals were intraperitoneally injected with OPP and back skin was collected 1-hour post-injection. The highest protein synthesis rates can be found in the inner root sheath (IRS) of the hair follicle (arrows). The lower right panel shows the quantification of OPP signal across basal and suprabasal epidermis, with a 1.75 times higher signal in suprabasal. N=33 quantifications across 2 sections from one animal, average ± standard deviation, quantified using ImageJ. (B) UMI count distribution, number of genes quantified and percent mitochondrial genes of single-cell RNA sequencing data in young and old skin. Clusters refer to Figure 3B. (C) Heatmap displaying the expression profiles of the top marker genes across the single-cell RNA sequencing of young and old epidermis. (D) Pseudotemporal ordering of the single-cell RNA sequencing data using Monocle outlines the differentiation trajectory from epidermal stem cells to the suprabasal clusters and granular layer. (E) A scatterplot displaying transcriptional and translational changes in aged versus young epidermal stem cells, generated using the Seurat pipeline. The overall distribution of gene expression changes resembles the results obtained from the glmGamPoi pipeline, as shown in Figure 4A. (F) *Krt10*-positive cells exhibit a higher number of detected genes in epidermal stem cells of clusters 0 and 5. p value indicates a Wilcoxon test. (G) *Krt5* and *Krt14* unique read counts are not elevated in *Krt10*-positive epidermal stem cells of clusters 0, 3 and 5 (G2M). The boxplot shows the distribution of *Krt5* and *Krt14* reads of single epidermal stem cells. p value indicates a Wilcoxon test. (H) A site codon occupancy without the start codon region (first 9 nucleotides) indicates similarly higher occupancy in glutamate (GAA, GAG) and aspartate (GAC, GAU) codons. To control for the strong start peak, A site codon occupancy was also analyzed separately without the start peak and two subsequent codons and compared to the A site codon occupancy plot presented in Figure 3I. (I) P site codon occupancy in old versus young epidermal cell clusters. (J) Scatterplot displaying transcriptional and translational changes in aged versus young upper hair follicle cells. (K) Ribosome profiling tracks of *Jun* highlight widespread translation in the 5′ UTR, pointing to potential uORF-mediated regulation of the main ORF. (L) Ligand-receptor analysis of young and aged epidermis employing CellChat reveals tissue-wide alterations of E-cadherin and semaphorin signaling, related to Figure 4Q-S. No semaphorin 4 signaling was detected in the aged epidermis. EpSC, epidermal stem cells. UHF, upper hair follicle. MP, macrophages. (M) *Jun* and *Jund*-controlled genes are enriched among the genes transcriptionally induced in aged epidermal stem cells. 742 transcripts induced in aged epidermal stem cells (single-cell RNA sequencing data) were analyzed in Enrichr using the ChEA (ChIP-X Enrichment Analysis) 2022 database.

